# A DNA-scaffolded Retron Ring Mediates Antiphage Immunity

**DOI:** 10.64898/2026.07.14.737667

**Authors:** Arturo Carabias, Ruiliang Zhao, Ricardo Garcia-Martin, Nuria Ardid-Muñoz, Mario Rodríguez Mestre, Huijuan Li, Johannes Anton Kühn, Josepha Magdalene Klas, Sarah Camara-Wilpert, Dennis Zhang, Nanna Wagner, Selma Margarita Kuypers, Tillmann Pape, Søren Johannes Sørensen, Rafael Pinilla-Redondo, Guillermo Montoya

## Abstract

Retrons are a diverse family of bacterial defence systems that couple reverse transcriptase (RT) activity to antiphage immunity by producing multicopy single-stranded DNA (msDNA). Yet, the molecular mechanisms by which most systems detect infection and execute immunity remain unknown. Here, we define the structure, immune sensing and effector mechanisms of retron type IX, composed of an RT, a non-coding RNA, a tandem KH-HTH subunit and a HEPN RNase effector. Retron IX confers robust protection against diverse bacteriophages. Cryo-EM and biochemical analyses reveal a supramolecular ring assembly composed of 8-10 repeating modules, spanning up to 230 Å in diameter. Within this assembly, the msDNA scaffolds the outer rim inhibiting the HEPN ribonuclease in a catalytically poised state, with the active site occluded within the inner layer. We identify a phage-encoded PD-(D/E)XK nuclease as the immune trigger that cleaves the exposed msDNA stem-loop, leading to ring disassembly and HEPN effector release. Upon activation, the HEPN ribonucleases cleave tRNAs indiscriminately, inducing dormancy and blocking phage propagation. Our findings unveil how ring-shaped higher-order assembly controls enzymatic activities while allowing signal-specific activation and establish the ncRNA as an evolvable scaffold underlying the remarkable structural and defensive diversity of bacterial retron immune complexes.

## INTRODUCTION

Retrons are bacterial reverse transcriptases (RTs) associated with non-coding RNAs (ncRNAs) that produce unusual DNA molecules known as multicopy single-stranded DNAs (msDNAs)^1^. The ncRNA folds into a conserved secondary structure containing msr and msd regions that binds the RT and prime reverse transcription. Priming occurs from the 2’-hydroxyl of a conserved guanosine within the *msr*, producing a chimeric DNA-RNA linked molecule through a unique 2’,5’-phosphodiester bond. The RNA template is subsequently degraded by host RNase H^2–4^, and in some cases the DNA-RNA hybrid is further processed by other host enzymes^5,6^.

The capacity of retrons to produce defined ssDNA molecules *in vivo* has been harnessed as a versatile tool for biotechnological applications, including genome editing^7–9^, high-throughput functional screenings^10^ and multiplexed genomic recording^11^. However, the biological role of retrons remained enigmatic for decades, until recent studies showed that they cluster with diverse toxic effectors^12^ to function as anti-phage defence systems operating through abortive infection^13–15^.

Mechanistic studies of diverse retron families (Type II^15–17^, Type I-A^18–21^, Type I-B2^22,23^ and Type I-C1^24,25^) have revealed that the RT-ncRNA-msDNA complex functions as an antitoxin, repressing cognate toxic effectors through direct binding. However, repression strategies vary markedly across families and are associated with distinct oligomeric states, ranging from relatively small assemblies^18–21,24,25^ to large supramolecular complexes^22,23^ and filamentous architectures^16,17^. During phage infection, phage-encoded proteins bind to or modify the msDNA, releasing effector inhibition and triggering cell death or dormancy to arrest infection^15–25^. Retron effectors are highly diverse^12^ and block phage replication through distinct mechanisms, including membrane-disruption^13,26^, depletion of essential metabolites^16,17^, and cleavage of nucleic acids^18–25^. Notably, recent evidence suggests that type VI retrons operate via a non-canonical mechanism in which phage infection induces msDNA production to activate translation of the toxic effector^26^, suggesting that additional novel retron molecular mechanisms remains to be discovered. Despite this rapid progress, the molecular mechanisms by which the remaining families sense phage infection and execute immunity remain poorly characterized.

Here, we define the structural and molecular mechanism of the retron IX family encoding a HEPN ribonuclease effector. We show for the first time that retron IX provides protection against diverse bacteriophages by inducing cell dormancy. We find that the RT, msrRNA, msDNA, HTH–KH protein, and HEPN effector assemble into a supramolecular ring-shaped complex of up to 230 Å in diameter. In this assembly, the msDNA acts as a structural scaffold that stabilizes the ring and restrains the HEPN ribonuclease in an inhibited state. Furthermore, we identify a phage-encoded nuclease as an immune trigger that cleaves the exposed msDNA stem-loop, disrupting the retron assembly and releasing HEPN activity. Upon activation, the HEPN effector cleaves host tRNAs at the anticodon-loop, leading to cell dormancy and blocking phage replication. These findings establish type IX retrons as novel supramolecular immune complexes and illustrate how higher-order assembly evolves to control the toxicity of diverse effectors.

## RESULTS

### Retron IX mediates defence against phages

Type IX loci are frequently carried by mobile genetic elements, including (pro)phages and plasmids (Figure S1A), and are enriched in genomic regions encoding known antiviral defence systems, consistent with a role in antiphage defence (Figure 1A). Type IX retrons are defined by a conserved operon organization encoding a reverse transcriptase (RT), a non-coding RNA (ncRNA), and two accessory proteins containing a helix-turn-helix (HTH) and an effector consisting of a HEPN domain (Figure 1A)^12^. The ncRNA is highly conserved and reliably identified by covariance models, displaying the canonical msr-msd architecture of retron elements, in which the msrRNA folds into a structured scaffold that supports RT binding and primes reverse transcription (Figure S1B). Homology-guided comparative genomic analyses show that type IX retrons are broadly distributed across bacteria, spanning 68 genera, with highest prevalence in Pseudomonadota (Figure S1A, Table S1).

**Figure 1.**
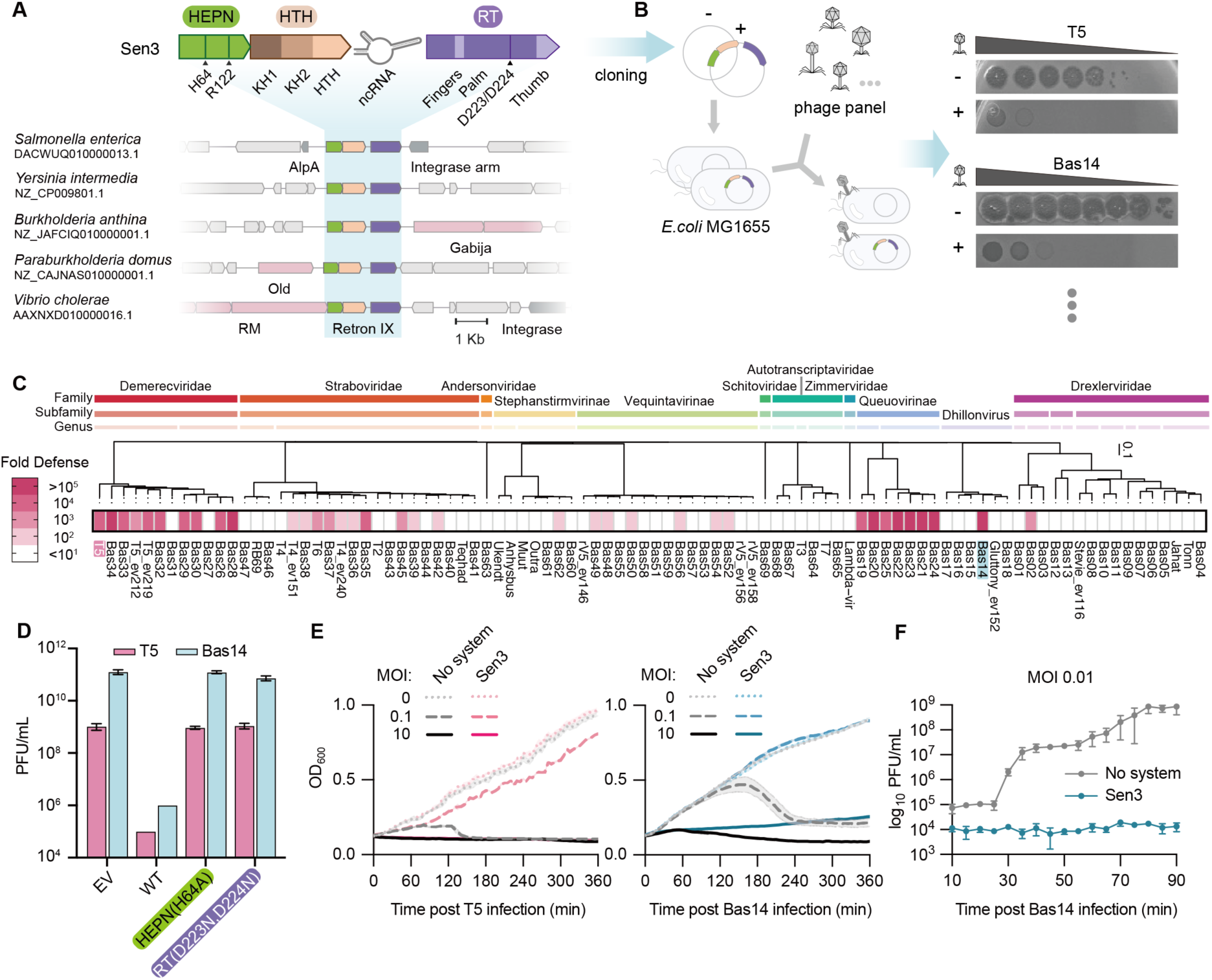
- Retron IX provides strong immunity against diverse phages. **A)** Top: Schematic representation of the operon organization and domain architecture of Retron-Sen3. Catalytic residues of the HEPN and reverse transcriptase (RT) domains are indicated with arrows. Bottom: Representative genomic contexts of retron IX systems, with known defence genes shown in pink and mobile genetic element-related genes in dim grey. **B)** Schematic of the phage infection assay and representative plaque assays showing reduced plaque formation of Sen3–expressing bacterial lawns relative to controls. **C)** Anti-phage activity of Sen3 against a panel of 92 coliphages, quantified as the fold change in efficiency of plating (EOP) relative to the empty vector control in *E. coli* MG1655. Data represent the mean of three biological replicates. **D)** Plaque-forming units (PFUs) of representative phages T5 and Bas14 infecting *E. coli* MG1655 carrying an empty vector, wild-type Sen3, or active-site mutants in the RT or HEPN domains. Data are means ± SD from three biological replicates. **E)** Growth of *E. coli* MG1655 carrying an empty vector or Sen3 after Bas14 and T5 infection at MOIs of 0.1 or 10. Data are means ± SD from three biological replicates. **F)** One-step growth curves showing Bas14 PFUs over time in *E. coli* MG1655 carrying an empty vector or Sen3. Data are means ± SD from three biological replicates.

To test their potential antiviral function, we selected a representative type IX retron from *Salmonella enterica* subsp. arizonae strain 2095-71/380, hereafter Retron-Sen3 (Sen3), cloned the intact operon under its native promoter into a low copy number plasmid, and expressed in *Escherichia coli* MG1655 (Figure 1B). Sen3 conferred broad and robust antiviral activity against a diverse panel of coliphages, with reductions in plaquing efficiency of up to five orders of magnitude against multiple phages (Figure 1C-D). Notably, mutations of conserved catalytic residues within the active site pockets of either the RT (D223N and D224N) or HEPN (H64A) abolished defence, demonstrating that the enzymatic activities of both RT and HEPN are required for anti-phage defence (Figure 1D, Figure S1C-E).

To determine how Sen3 protects bacteria from phage infection, we monitored bacterial culture growth during infection across a range of multiplicities of infection (MOI). At low MOI, Sen3-expressing cultures grew comparably to uninfected controls, whereas cells lacking the system succumbed to infection. In contrast, at high MOI, both Sen3-expressing and control cultures exhibited growth collapse (Figure 1E), consistent with a population-level defence mechanism. To directly quantify the impact on phage propagation, we measured phage progeny production during single-cycle infections. In control cultures, infection produced an average burst size of ∼100 progeny phages per infected cell within ∼40 min, whereas no detectable phage progeny was produced in cells expressing Sen3 (Figure 1F). These dynamics are consistent with abortive infection, in which infected cells undergo suicide or growth arrest to contain phage epidemics. Together, these results establish Sen3 as a broadly acting antiviral defence system amenable to mechanistic dissection, raising questions about how phage infection is sensed and coupled to effector activation.

### Assembly of a retron IX system defence complex

Retron complexes typically rely on protein-protein interactions that can be reliably predicted by AlphaFold3^27^ (AF3). Accordingly, AF3 predicted with high confidence a ternary Sen3 complex in which the HTH domain bridges the RT and HEPN domains (Figure S2A-C). N-terminal to the HTH domain, a tandem KH module was also identified, although it did not engage in interactions within the ternary complex model. To test these predictions, we introduced point mutations at key interfacial residues and assessed anti-phage defence. All mutations abolished immunity, confirming the functional importance of these contacts (Figure S2D).

To gain structural insight into the retron assembly, we overexpressed Sen3 with an inactivating HEPN-domain mutation (H64A) to reduce potential toxicity in a modified *E. coli* MG1655-DE3 strain^16^ and attempted purification. However, the HEPN effector remained insoluble after lysis, precluding further *in vitro* characterization (Figure S3A and S3B). To overcome this limitation, we searched for retron IX homologs sharing the architectural features of Sen3, including conserved domain organization, AF3-predicted interfaces, catalytic residues, and retron ncRNA features, and identified a retron from *Yersinia intermedia*, hereafter Retron-Yin1 (Yin1), with a predicted higher HEPN domain solubility, as assessed by CamSol^28^ (Figure S1C-E and S3C-H). Expression and purification of Yin1 yielded soluble HEPN and a stable complex comprising all three proteins together with msrRNA and msDNA (Figure S3C, S3I and S3J; STAR Methods), which was suitable for structural determination.

### Cryo-EM reveals a supramolecular ring architecture of Retron-Yin1

We next used single-particle cryogenic electron microscopy (cryo-EM) to image Retron-Yin1 at different sample concentrations, obtaining maps at 3.24 Å and 7.20 Å resolution (Figure S4, S5; Table S2; STAR Methods). We observed ring-shaped supramolecular assemblies in the micrographs corresponding to Yin1 assemblies at low sample concentration (Figure 2A-D, S4A, S5A-F; STAR Methods). However, strong preferential orientation of the rings precluded accurate map interpretation under these conditions (Figure S5F). In contrast, the rings were partially disrupted at higher concentration, improving particle distribution and reducing preferential orientation, which produced higher-quality maps used for model building (Figure S4B, S5G-L; STAR Methods).

**Figure 2.**
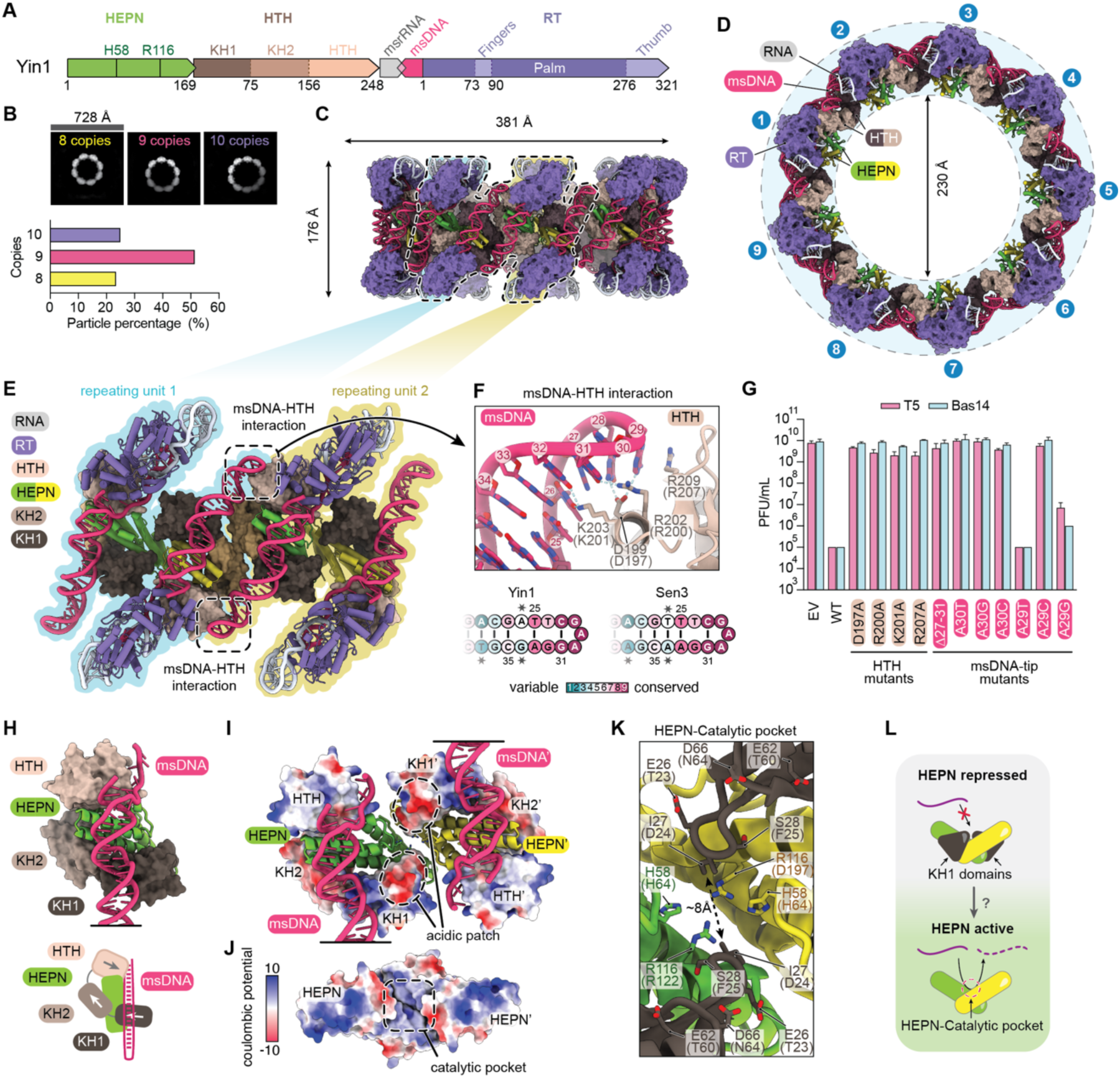
- Structure of the Retron-Yin1. **A)** Domain architecture of the Retron-Yin1. **B)** Representative 2D class averages of Yin1 rings displaying 8-10 copies, with quantification. **C)** Side and **D)** top views of the Yin1 structure displaying a nonameric assembly. **E**) Assembly of two repeating units highlighting the interactions between the msDNA-tips and the HTH proteins **F)** Close-up view of the interaction between the msDNA-tip and conserved residues in the HTH domain. Residue numbering corresponds to Yin1, with homologous Sen3 residues indicated in parentheses (see. Figure S1C-E). Evolutionary conservation of the msDNA tip in Yin1 and Sen3 retrons is shown schematically bellow as a cartoon. **G)** Plaque-forming units (PFUs) of T5 and Bas14 phages infecting *E. coli* MG1655 strains carrying an empty vector or expressing wild-type, msDNA mutants, or HTH mutants. Data represent mean ± SD from three biological replicates. **H)** View of the KH1, KH2, and HTH domains interacting with the HEPN domain locked by the msDNA. **I)** View of the HEPN dimer within the complex, with electrostatic surface potential mapped onto KH1, KH2, and HTH domains. **J)** Surface representation of the HEPN domain highlighting its electrostatic potential and the basic patch harboring the catalytic residues. K) Close-up view of the HEPN catalytic pocket, showing catalytic residues and negatively charged KH1 residues as sticks. Residue numbering corresponds to Yin1, with homologous Sen3 residues in parentheses. **L)** Schematic representation of HEPN inhibition mediated by the KH1 domains.

The msDNA forms the outer scaffold of the rings (Figure 2C and 2D), while the HEPN domains occupy the inner layer, where they form ribonuclease dimers (Figure S6A). These dimers constitute a catalytically competent conformation in which conserved residues from both protomers contribute to a composite active site^29^ (Figure S6B). The rings are composed of 8-10 repeating units, with 9-mers being the most abundant class (∼50%) (Figure 2B). Each unit comprises a HEPN dimer, with each protomer bound to an RT/HTH-KH/ncRNA/cDNA module (Figure S6C). This arrangement closely matches AF3 predictions for Yin1 and agrees with Sen3 models (Figure S6D), supporting a conserved architecture across type IX retrons. A closer inspection of the interfaces revealed that ring formation is mediated by lateral interactions between adjacent units, involving contacts between msDNA stem-loop tips and the HTH domain of neighbouring units (Figure 2E and 2F). Four conserved HTH residues D199, R202, K203 and R209 (corresponding to D197, R200, K201 and R207 in Sen3), associate with the msDNA (Figure 2F and S1E). D199 coordinates dG28 and dA29, R202 contacts dG31, and K203 coordinates dT26 and dG32, mediating sequence-specific recognition, while R209 stacks against dA29 (Figure 2F). Mutation of these residues in Sen3 abolished defence, phenocopying deletion of the msDNA tip (Δ27–31) (Figure 2F and 2G). Additionally, nucleotide substitution analysis showed that dA30 is essential, whereas dA29 tolerates limited variation (Figure 2G).

### msDNA-KH architecture inhibits the HEPN subunit

The position of the msDNA further stabilizes the tandem KH domains in a conformation that wraps around the HEPN dimer (Figure 2H, Figure S6C and S6E-F). In this engaged state, conserved complementary interfaces between KH and HEPN domains are observed in both Yin1 and Sen3 retrons (Figure S6F-G), reinforcing a shared structural organization. Notably, the KH1 domains bind atop each HEPN protomer, masking a positively charged surface that provides access to the catalytic site (Figure 2I-K). This arrangement represents an inhibited state, restricting the electrostatic interactions required for substrate binding (Figure 2L), which have been recently observed in the HEPN domain of the structure of a Cas13-tRNA complex^30^. Access to the active site is further limited by the negatively charged msDNA backbone (Figure 2I), an acidic patch on KH1 (Figure 2I), and two hydrophobic residues (I27) located at the edges of the slit formed by the two KH1 domains (Figure 2K). Collectively, these observations suggest that, during phage infection, the HEPN domain may be released from this inhibited conformation, thereby enabling its activation.

### A phage-encoded nuclease triggers immunity

Phages often escape antiviral defences through mutations in factors that trigger immunity, a principle that has inspired experimental evolution approaches to identify phage-encoded determinants of immune activation^31^. To investigate how retron type IX is activated, we isolated phage escapers that overcome Sen3-mediated defence and mapped the underlying mutations by whole-genome sequencing and comparative genomics relative to the corresponding wild-type phages (Figure 3A). We focused on phages from the Queuovirinae subfamily (Bas19–Bas25), which were uniformly sensitive to Sen3 (Figure 1C). However, we failed to isolate escaper mutants, suggesting that the triggering factor might be essential for phage viability. In contrast, isolation and sequencing of Bas14 escapers revealed mutations mapping to a single locus, Bas14_0045, hereafter B14T. These included missense mutations and premature protein truncations (Figure 3B and S7A), strongly implicating this gene product as a determinant of retron IX sensitivity.

**Figure 3.**
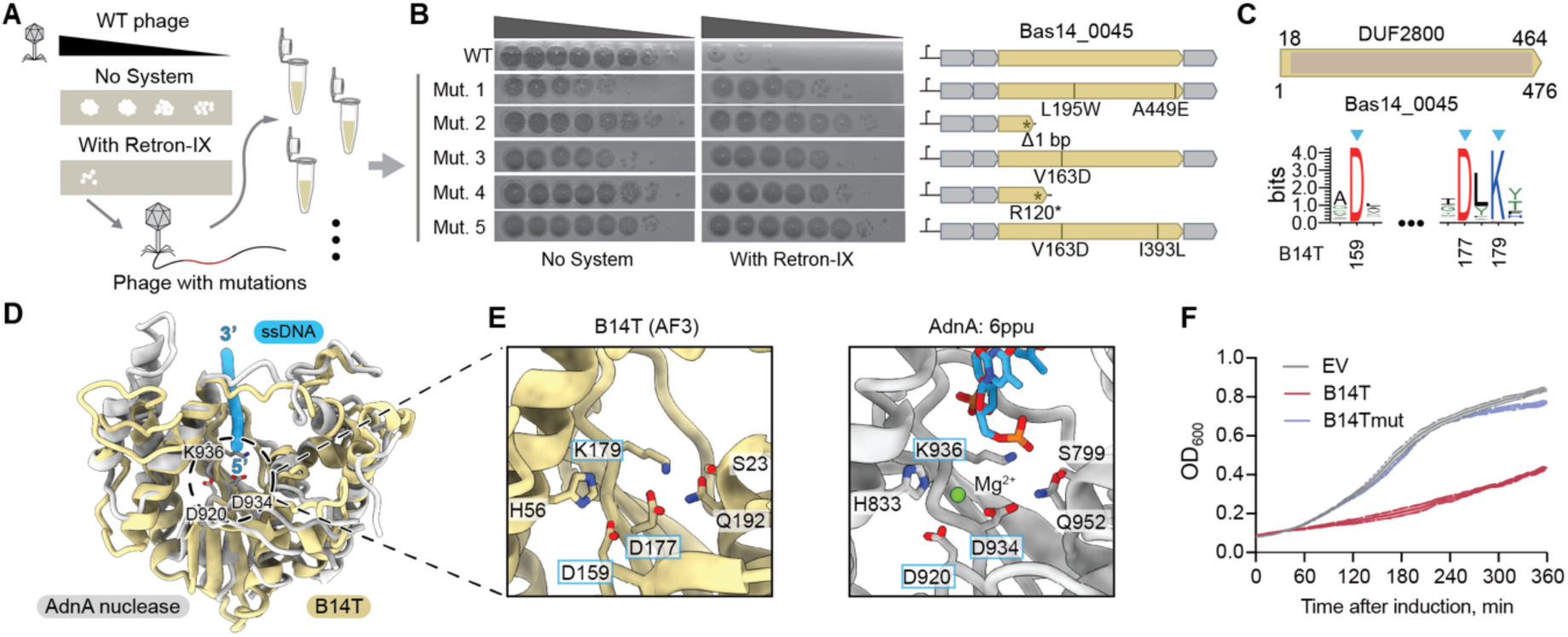
- A phage nuclease triggers retron IX anti-phage defence. **A)** Schematic of the escaper isolation approach to identify phage-encoded triggers of Sen3. **B)** Bas14 escaper phages carrying mutations in Bas14_0045 evade retron-mediated defence. Plaque assays were performed with wild-type (WT) Bas14 and escaper mutant phages (Mut. 1–5) on *E. coli* expressing either an empty vector (No System) or Sen3. **C)** Domain organization of Bas14_0045, with a sequence logo showing conservation of the PD-(D/E)XK motif among phage-encoded homologs. Conserved catalytic residues are shown marked with blue arrows. **D)** Structural comparison of the AF3-predicted Bas14_0045 trigger protein (B14T) (pTM=0.66) with the end-resection nuclease AdnA (PDB: 6PPU; RMSD: 9.99 Å across 218 pairs). **E)** Zoom of the active site showing residues in B14T (AF3) and AdnA (PDB: 6PPU). Residues conforming the PD(D/E)XK motif are labelled with a blue frame. **F)** Toxicity testing of the B14T trigger co-expressed with Sen3. Data are means ± SD from three biological replicates.

B14T is a protein of unknown function comprising a DUF2800 domain (Figure 3C, and S7A-B). Structural comparison analyses revealed a conserved catalytic domain closely resembling that of AdnA and RecB DNA-resection nucleases, bearing a canonical PD-(D/E)XK motif, followed by a predicted C-terminal disordered region (Figure 3D-E, Figure S7C-F). To test whether this phage protein can trigger retron IX, we co-expressed B14T from a plasmid together with Sen3 and monitored bacterial culture growth in the absence of phage infection (Figure 3F). This resulted in pronounced growth inhibition, indicating that B14T is sufficient to trigger Sen3-mediated toxicity. Consistent with this, mutation of residues within the predicted catalytic pocket (D177A/K179A) abolished toxicity (Figure 3E and 3F). Taxonomic analysis of B14T homologs showed that they are broadly distributed across diverse phages and are associated with hosts spanning diverse bacterial phyla (Figure S7G). This distribution is consistent with the broad anti-phage activity of retron IX (Figure 1C; Table S1). Collectively, these results identify B14T as a phage-encoded activator of Retron-Sen3 and demonstrate that its nuclease activity is required for immune activation.

### B14T cleaves the base of the msDNA stem-loop

The msDNAs lock the KH domains in a conformation that shields the HEPN catalytic site within the ring assembly, suggesting direct control of HEPN activity by the msDNA (Figure 2H and 2I). We therefore hypothesized that the phage-encoded nuclease B14T triggers retron immunity by cleaving the msDNA. Co-expression of wild-type B14T with Sen3 expressed from its endogenous promoter led to a complete loss of the msDNA band, whereas mutation of the B14T catalytic pocket (D177A/K179A) abolished this effect (Figure 4A), thereby supporting that msDNA cleavage depends on B14T nuclease activity.

**Figure 4.**
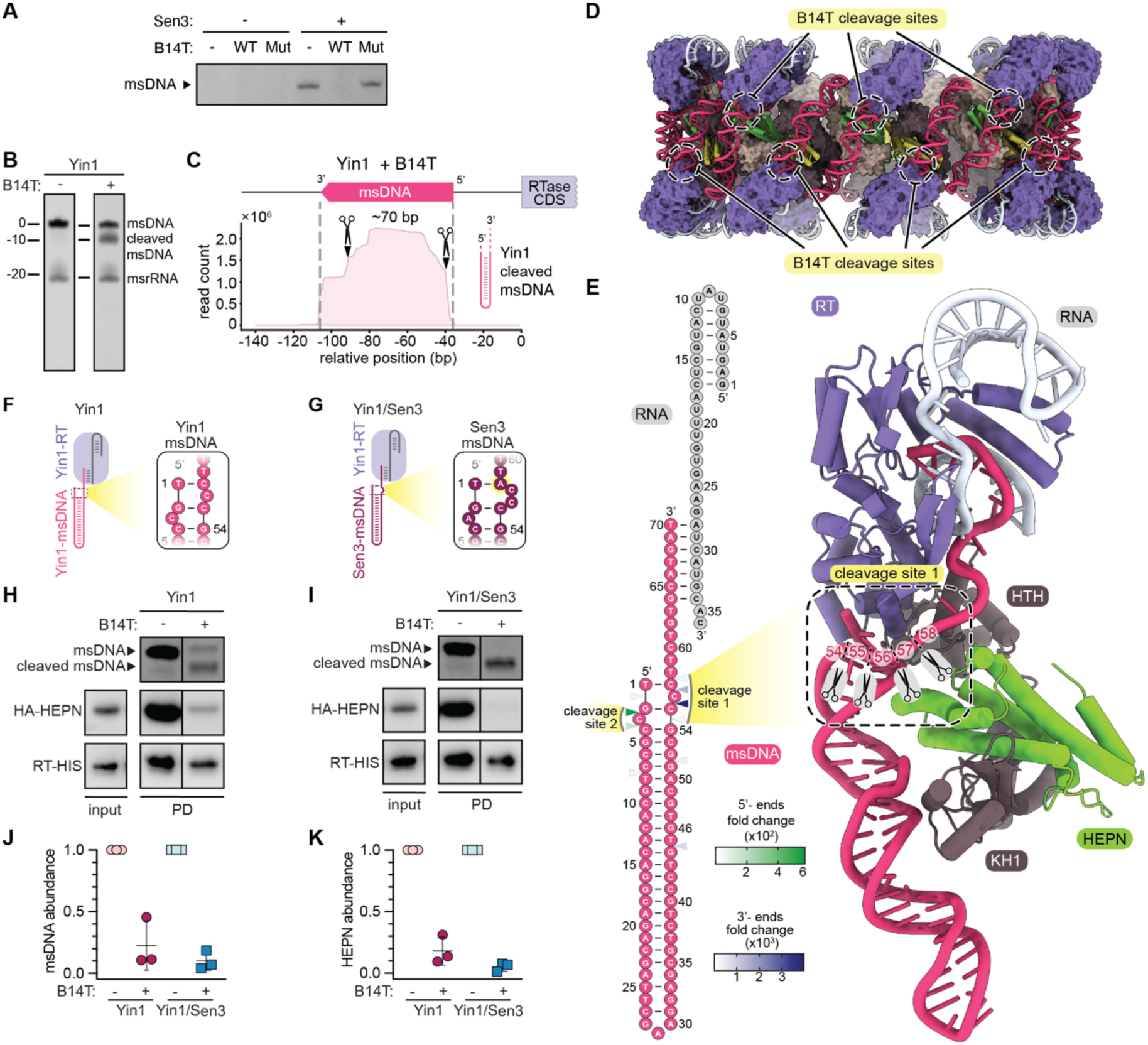
- The phage nuclease cleaves retron msDNA and releases the HEPN domain. **A)** TBE-urea gel showing Sen3 msDNA upon co-expression with B14T wild-type (WT) or catalytic inactive mutant (D177A/K179A). Representative experiment of three biological replicates. **B)** *In vitro* cleavage of Yin1 msDNA by the B14T catalytic domain following 15 min incubation (see Figure S3A). The image is representative of more than three independent experiments. **C)** Read counts of Yin1 msDNA after incubation with purified B14T catalytic domain, determined by next-generation sequencing. The two major cleavage sites are indicated. **D)** Side view of Yin1 assembly highlighting the position of B14T cleavage sites at the base of the msDNA stem-loops. **E)** Schematic representation (left) and atomic model of Yin1 msrRNA and msDNA (right). The two B14T cleavage sites are highlighted in yellow. **F-G)** Schematic representation of Yin1 (F) and (G) a Yin1/Sen3 chimera, with close-up views of cleavage sites 1 and 2. The additional nucleotide present in Sen3 is highlighted in yellow. **H-i)** Pull-down assays of Yin1 (H) and the Yin1/Sen3 (I) chimera before and after incubation with B14T lysates. Representative experiment of three biological replicates. **J)** Quantification of msDNA abundance in (H, I), calculated as the signal of the non-cleaved band (upper band) normalized to RT levels and per retron type. **K)** Quantification of HEPN abundance in (H, I), measured as the chemiluminescent signal normalized to RT levels and per retron type.

To test whether B14T directly cleaves retron msDNA, we incubated purified Yin1 with purified B14T comprising the catalytic domain (STAR Methods). This resulted in msDNA cleavage and the appearance of a product approximately 10 nt shorter than full-length msDNA (Figure 4B, Figure S8A). Next-generation sequencing identified two predominant cleavage sites on complementary strands at the base of the msDNA stem-loop (Figure 4C-E, Figure S8B-D). Cleavage site 1 spans nucleotides dG54-dT58, with the phosphodiester bond between dC55 and dC56 most frequently cleaved, while cleavage site 2 encompasses nucleotides dT1-dC4, with preferential cleavage between dG2 and dC3. In the structure, the msDNA coats the outer surface of the ring (Figure 4D), where site 1 is solvent-exposed whereas site 2 is partially occluded by the HTH and HEPN domains (Figure S8C-D). This suggests that initial cleavage at site 1 may expose site 2 for subsequent cutting.

### msDNA cleavage unlocks the HEPN effector

As the base of the msDNA stem-loop anchors the DNA to the RT, we reasoned that complete cleavage would release msDNA from the complex. Consistent with this notion, mixing lysates expressing B14T with Yin1 followed by His-tag pull-down of the RT led to a marked reduction in msDNA abundance (Figure 4F-J). Despite the high sequence conservation of msDNA between the two homologous retrons, we noted that Sen3 msDNA is 71 nt long, whereas Yin1 msDNA is 70 nt (Figure 4E and S8E). The additional nucleotide in Sen3 is positioned within the primary B14T cleavage site (Figure 4E and S8E). To determine whether this difference influences cleavage efficiency, we generated a chimeric Yin1 in which the msDNA stem-loop and single-stranded DNA region were replaced by those of Sen3 (Yin1/Sen3; Figure 4F-G). B14T cleaved this chimera more efficiently than wild-type Yin1, indicating that the unpaired nucleotides in this region increase susceptibility to B14T-mediated cleavage (Figure 4H-J).

Notably, msDNA cleavage was accompanied by a marked reduction in HEPN abundance within the complex, consistent with dissociation of HEPN from the RT-HTH module (Figure 4H-K). Structural comparison with an AF3 model generated in the absence of msDNA suggests that this dissociation may be driven by a reorientation of the KH domains (Figure. S8H-I). The msDNA–HTH interaction also constitutes the only bridge between adjacent subunits in the ring (Figure 2E-F, Figure 4D). Thus, cleavage of the msDNA disrupts the structural continuity required to maintain the higher-order assembly, likely promoting ring disassembly. Together, these data support a model in which msDNA cleavage destabilizes the ring and promotes HEPN dissociation.

### HEPN effector degrades RNA upon phage-triggered activation

HEPN domains are conserved across prokaryotes and eukaryotic organisms, functioning as metal-independent RNases^32,33^. In bacterial defence systems, HEPN domains can act as effectors that degrade RNA to restrict phage propagation ^19–21,24,34–37^. We therefore hypothesized that retron IX systems mediate antiphage defence through cleavage of cellular RNAs. To test this, we performed urea-PAGE analysis of nucleic acids extracted from cells co-expressing Sen3 and B14T (Figure 5A-B, Figure S9A). This revealed a marked accumulation of small RNA fragments (∼30-50 nt) and a concomitant reduction in RNA species of ∼70-100 nt. To dissect the underlying mechanism, we combined wild-type or catalytically inactive variants of B14T and the HEPN effector. The cleavage products were detected only when both B14T and the HEPN effector were catalytically active, consistent with RNA cleavage upon Sen3 activation (Figure 5B).

**Figure 5.**
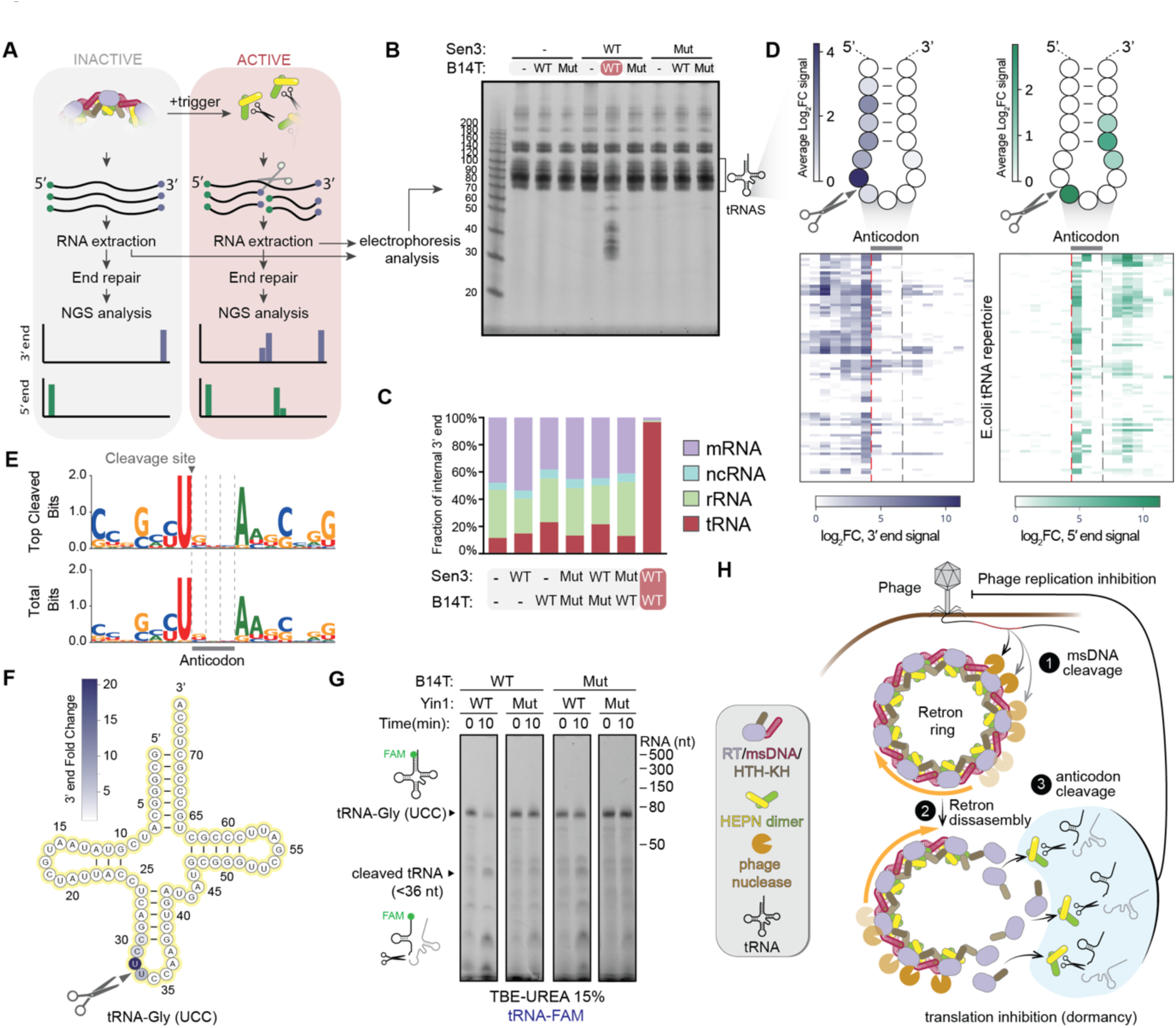
- Retron IX cleaves tRNAs at the anticodon-loop. **A)** Schematic of the RNA-seq approach used to assess *in vivo* RNA cleavage activity. **B)** Urea-PAGE analysis of small RNAs (<200 nt) extracted after trigger gene induction from cells harbouring the indicated empty vector control, active, or inactive constructs for Sen3 and B14T (Sen3 Mut: HEPN H64A; B14T Mut: D177A/K179A). The gel is representative of three biological replicates. **C)** Fraction of internal 3′ ends in different annotated RNA types for selected samples from the experiment shown in (b). **D)** Schematic depiction of the tRNA anticodon-loop and position-specific signal across the *E. coli* tRNA repertoire. Circles represent tRNA nucleotides and are coloured by the average log₂ fold change in 3′-end (left) or 5′-end (right) CPM signal between Sen3-B14T and Sen3_Mut-B14T samples. Heatmaps show the corresponding signals for individual annotated E. *coli* tRNAs. Sequences include the anticodon and seven nucleotides upstream and downstream. **E)** Sequence logos of the anticodon-loop region for tRNAs showing >30-fold change and for all tRNAs. Sequences include the anticodon and seven nucleotides upstream and downstream. **F)** Schematic secondary structure of tRNA-Gly(UCC), with each nucleotide coloured according to the fold change in 3′-end CPM signals between Sen3-B14T and Sen3_Mut–B14T samples. **G)** *In vitro* tRNA cleavage assay with non-activated and activated Yin1 wild-type and inactive mutant. FAM-labelled tRNA-Gly(UCC) was used as a substrate for cleavage. The gel is representative of two independent experiments. **H)** Proposed model of the retron IX system immune mechanism. Yin1 forms a ring-shaped supramolecular complex that inhibits the HEPN nuclease. During phage infection, a phage-encoded nuclease cleaves msDNA (1), triggering disassembly of the retron complex and release of HEPN (2). HEPN then cleaves tRNAs to inhibit translation and restrict phage propagation (3).

### The HEPN cleaves the anticodon-loop of tRNAs

RNA cleavage generates products with discrete 5′ and 3′ read termini that can be mapped by small RNA sequencing (Figure 5A). To identify HEPN-mediated cleavage sites, we performed small RNA sequencing after end-repairing (<200 nt) on Sen3-B14T-expressing cells 60 min after induction. Compared with catalytic mutant and empty vector controls, internal RNA ends were significantly enriched at tRNA loci in cells expressing wild-type Sen3-B14T, identifying tRNAs as the primary cellular targets of Sen3 (Figure 5C, Figure S9B).

To define the cleavage site, we analysed read termini across all annotated tRNAs. Enriched 3′ ends mapped predominantly to the nucleotide immediately upstream of the anticodon and enriched 5′ ends to the first anticodon nucleotide, together pinpointing cleavage to the first anticodon position (Figure 5D, 5F and S9C). Additional enriched ends likely reflect secondary cleavage events or host RNA trimming^38,39^. All read ends enriched over 30-fold upon Sen3 activation mapped exclusively to tRNA, supporting tRNAs as the predominant targets (Table S3). Comparison of anticodon-loop sequences across the complete *E. coli* tRNA repertoire revealed no consensus cleavage motif, suggesting that substrate recognition is governed by anticodon-loop structure rather than sequence (Figure 5E).

To test whether the HEPN domain directly cleaves tRNAs, we reconstituted Yin1 activation *in vitro* by incubation with the phage trigger B14T and assayed its RNase activity on a synthetic FAM-labelled tRNA-Gly(UCC). We included a catalytically inactive B14T mutant (D177A/K179A) to control for activation-independent RNA cleavage, and a HEPN-inactive Yin1 mutant (R116A) to confirm that cleavage is HEPN-dependent. Activated Yin1 cleaved the tRNA, while this activity was not apparent in the control samples (Figure 5G, S9E and S9F). Remarkably, a ∼36-nt cleavage product was detected, as estimated by comparison with the msrRNA band (Figure S9E and S9F), consistent with cleavage at the anticodon-loop. Altogether, these results demonstrate that the phage B14T nuclease triggers activation, deploying a HEPN-domain endoribonuclease that depletes host tRNA pools through anticodon-loop cleavage to mediate antiphage defence.

## DISCUSSION

Here, we show that Retron-Yin1 assembles into a closed, symmetric supramolecular ring comprising 8-10 protomers (Figure 2). In this architecture, the msDNA serves both as the structural “glue” that stabilizes the assembly and as a regulator of the HEPN effector. Specifically, the msDNA positions the KH domains over the HEPN active site, thereby inhibiting its activity by blocking substrate access (Figure 2H-L). These findings reinforce the known roles of msDNA beyond immune sensing, as a structural regulator of enzymatic activity^15–25^. The HTH-KH and HEPN proteins represent the hallmark feature of type IX retrons. Although KH domains are classically associated with RNA binding^40^, here they appear to serve primarily structural and regulatory roles. This suggests that this new role of the KH domain might have originated from an ancestral state in which the KH domain cooperated with the HEPN effector in RNA recognition.

We identified a phage-encoded nuclease, B14T, carrying a conserved PD-(D/E)XK motif as the trigger of Sen3. Thus, B14T joins the catalogue of phage-encoded msDNA-interacting proteins that activate retrons, including nucleases^15,18,20,21,24,25^, methylases^15,17^, and single-stranded DNA binding (SSB) proteins^22,23^. These observations reinforce the emerging notion that the msDNAs in retrons act as molecular decoys of the natural substrates of phage-encoded DNA-interacting proteins, and that their modifications act as signals to activate immunity. Although the function of B14T in the phage life cycle remains unknown, PD-(D/E)XK endonucleases are broadly implicated in restriction-modification systems, DNA repair, and recombination^41^, suggesting that B14T may contribute to phage genome processing or recombination. Notably, several defence systems appear to be triggered by phage-encoded DNA repair and recombination machineries^26,31^, raising the question of why this convergence exists. As recombination modules help phages and plasmids repair breaks caused by genome-targeting systems such as restriction-modification and CRISPR-Cas^42–44^, sensing these repair modules may allow bacteria to detect mobile genetic elements that are pre-armed to escape genome-targeting defences. This may explain the frequent co-localization of type IX retrons with DNA-targeting systems in defence islands (Figure 1A), which would be expected to functionally synergize against such phages. Together, these layered defence architectures may create a double bind for phages, in which the very mechanisms that enable escape from one defence system leads to the activation of another.

Activated Sen3 and Yin1 cleave tRNAs, placing type IX retrons within a growing family of antiviral defence systems that exploit tRNA depletion to block viral propagation (Figure 5A-G). This includes, CRISPR-Cas13^35^, the PARIS system^37^, Schlafens^34^, Type I-A retrons (e.g., Eco7) ^19–21,36^ and Type I-C1 retrons (e.g., Eco2)^24^. In contrast with these systems, type IX Sen3 retron HEPN cleaves tRNAs indiscriminately in a sequence-independent manner, suggesting that anticodon-loop shape rather than sequence determines substrate selection. (Figure 5D-F). Phages that evade tRNA-targeting defence systems frequently encode tRNA genes with mutations that render them resistant to host anticodon nucleases^36,45^. In this context, the broad tRNA targeting activity of the HEPN effector suggests that it not only ensures rapid shutdown of protein synthesis but simultaneously prevents phages from escaping through the acquisition of resistant tRNA variants.

Retron systems build immune complexes using nucleic acid scaffolds. The high tolerance in sequence variation while preserving msrRNA and msDNA structure enables rapid evolution and likely facilitates the recruitment of novel toxic effectors, driving immune innovation and retron diversity. On a wider perspective, the formation of supramolecular assemblies by immune components is a principle conserved across all domains of life, and such higher-order architectures are thought to offer several evolutionary advantages in the context of immunity^46^. Notably, whereas immune systems that form ring-like structures at the membrane are widespread^47–49^, cytoplasmic ring assemblies are exceedingly rare in the bacterial immune repertoire. Yin1 and the recently described RAZR^50^ system together define this emerging class. Unlike RAZR, which senses and assembles around fixed-diameter phage protein complexes, Yin1 self-assembles with self-synthesised msDNA and exhibits heterogeneous multimeric states (typically 8-10 subunits). Remarkably, although both systems deploy HEPN effectors, their activation occurs through opposing mechanisms, with ring stabilization triggering RAZR and ring disassembly activating Yin1.

Collectively, our findings define type IX retrons as a novel class of supramolecular immune complexes that build unusual cytoplasmic ring architectures. These findings reinforce the msDNA as a dual-function element that both inhibits effector activity and senses phage infection cues, highlighting DNA synthesis as a versatile platform for immune innovation and enabling the engineering of retron-based biotechnologies.

## ACKNOWLEDGMENTS

We thank Alexander Harms for providing the BASEL phage collection and Lars Hestbjerg Hansen, Claudia Igler, Lone Brøndsted and Felix d’Hérelle Reference Center for Bacterial Viruses from the Université Laval for providing additional phages used in this work. We also thank Alejandro Gonzalez Delgado for suggestions on the msDNA sequencing; Inga Songailiene for suggestions on the RNA sequencing; Farooq Gulomar for the protein production support; Blanca López Méndez for mass spectrometry analysis; Andrew Juul Ramírez Mountain for the experimental support; the Danish cryo-EM National Facility at the University of Copenhagen for support during data collection supported by grant NNF14CC0001, NNF24SA0098829 to the Novo Nordisk Foundation Center for Protein Research (CPR). G.M. is part of CPR, which is supported financially by the Novo Nordisk Foundation (NNF14CC0001, NNF24SA0098829). This work was also supported by the ERC-AdG 101096548 (INTETOOLS), NNF0024386, NNF17SA0030214, and NNF18OC0055061 grants to G.M, who is a member of the Integrative Structural Biology Cluster (ISBUC) at the University of Copenhagen. A.C is supported by a Lundbeck Fonden grant R380-2021-1448. R.P.-R is supported by the Lundbeck Fonden grant R347-2020-2346 and VILLUM FONDEN VIL60763. R.Z. is supported by CSC Scholarship 202206760017.

## AUTHOR CONTRIBUTIONS

Conceptualization: A.C., R.Z., R.P.R., G.M.; Methodology: A.C., R.Z., R.G.M., N.A.M.; Investigation: A.C., R.Z., R.G.M., N.A.M, M.R.M, H.J, J.A.K, J.M.K, S.C-W, D.Z, N.W, S.M.K., T.P.,; Visualization: A.C., R.Z.; Writing – original draft: A.C., R.Z. R.P.R., G.M.; Writing – review & editing: A.C., R.Z., R.P.R., G.M.; Funding acquisition: A.C., S.J.S., R.P.R., G.M.; Supervision: A.C., R.P.R., G.M.

## SUPPLEMENTAL INFORMATION

**Figures S1-9**

**Table S1.- Taxonomy information of retron IX RT**

**Table S2.- Cryo-EM data collection and model refinement statistics**

**Table S3.- *In vivo* targets detected as cleaved by retron IX**

**Table S4.-Plasmids, oligonucleotides and phages**

## STAR METHODS

### RESOURCE AVAILABILITY

#### Lead contact

Further information and requests for resources and reagents should be directed to and will be fulfilled by the lead contact, Guillermo Montoya.

#### Materials availability

All reagents generated in this study are available upon request to the lead contact with a completed Materials Transfer Agreement.

#### Data and code availability

- The structure’s atomic coordinates and cryoEM maps have been deposited in the PDB (https://www.rcsb.org) and EMDB (https://www.ebi.ac.uk/emdb/) databases. Accession numbers and DOIs are listed in Table S2. Uncropped imaging data will be deposited at Mendeley Data (https://data.mendeley.com/).
- This paper original code will be included for revision.
- Any additional information required to reanalyze the data reported in this paper is available from the lead contact upon request.

### EXPERIMENTAL MODEL AND STUDY PARTICIPANT DETAILS

#### Bacterial strains and phages

*E. coli* NEB5α and *E. coli* MG1655 were used for plasmid construction and as hosts for phage-related experiments, respectively. NEB5α *was* grown in lysogeny broth (LB) medium at 37°C shaking at 180 rotations per minute (rpm). The strains used for phage experiments were grown in LB supplemented with 10 mM MgCl_2_ at 37°C shaking at 180 rpm. When appropriate, the media were supplemented with kanamycin (50 μg/mL), spectinomycin (2 μg/mL), ampicillin (100 μg/mL) and/or chloramphenicol (25 μg/mL) to maintain the plasmids. All phages used in this study are listed in Table S4.

### METHODS DETAILS

#### Plasmid construction

The plasmids vectors used in this study and the constructed plasmids are listed in Table S4. To express the proteins of interest (Table S4), their coding sequences, sometimes together with their native promoters as indicated, were amplified from phage or synthesized from Twist Bioscience. In general, the plasmids were constructed with ClonExpress II One Step Cloning Kit (Vazyme), USER cloning (NEB) or Infusion cloning kit (Takara). Site-directed mutagenesis was performed by polymerase chain reaction (PCR) amplification of the plasmids carrying the intact genes using the primers containing mutant sites, followed by assembly of the resulting plasmid fragments in *E. coli* NEB5α or Stellar competent cells (Takara). The primers used in the study are listed in Table S4.

#### Phage propagation and plaque assay

*E. coli* MG1655 was used as the host for phage propagation. Overnight cultures of *E. coli* MG1655 were diluted 1:100 in LB supplemented with 10 mM MgCl2 and grown to an OD_600_ of ∼0.2 at 37°C. Then the culture was infected with phages of interest at a multiplicity of infection (MOI) of 0.01 and further grown overnight. To separate the virions from cell debris, the cultures were centrifuged at 7000 g for 10 min and the supernatants were filtered through 0.22 μM membrane filters (Fisherbrand). The resulting phage shocks were stored at 4°C. The phage titre was ascertained by pipetting 10 μL droplets of serial dilutions onto a LBA overlay (0.5% w/v) supplemented with 10 mM MgCl_2_ that was seeded with 1% of host overnight culture. After an overnight incubation, plaques were enumerated, with the phage titre presented as plaque-forming units per milliliter (PFU/mL).

For plaque assays, *E. coli* host cells expressing an empty vector (pTU175) or wild-type or mutated retron IX systems were grown in LB medium supplemented with spectinomycin at 37°C overnight. Then 1% of the overnight culture was mixed with molten LB agar overlay (0.5% w/v) supplemented with the appropriate antibiotic and 10 mM MgCl₂, and the mixture was poured onto LB agar plates supplemented with 10 mM MgCl₂. Meanwhile, phages of interest were 10-fold serially diluted and 5 µL of the serial dilution were spotted on the bacterial overlay. The plates were then incubated overnight, and the plaques were counted to calculate the phage titer in PFU/mL. The fold defence was determined by dividing the plating efficiency (in PFU/mL) of the empty vector by the efficiency of plating on *E. coli* MG1655 carrying wild-type Sen3. Data are shown as the mean ± standard deviation (SD) of three biological replicates.

#### Phage-infection in liquid medium

Overnight cultures of *E. coli* MG1655 containing an empty vector or the Retron-Sen3 were diluted 1:100 into fresh LB medium supplemented with spectinomycin (2μg/mL) and grown to an OD₆₀₀ of ∼0.2 at 37°C. Then 180 μL of the cultures were transferred into the wells of a 96-well plate and 20 μL of the diluted phages were added into the cultures for a final MOI of 0.1 or 10, or 20 μL of LB media for the uninfected control. The plate was incubated at 37°C with shaking at 200 rpm in a BioTek Synergy H1 microplate reader, and OD₆₀₀ was measured every 5 min.

#### Phage burst assay

*E. coli* MG1655 cells carrying either the empty vector or the Retron-Sen3 were inoculated from overnight cultures into fresh LB medium at a 1% dilution and grown with shaking at 37°C to an OD₆₀₀ of ∼0.2. Phages were added at an MOI of 0.01 based on the calculated PFU/mL, and the mixtures were incubated statically at 37°C for 5–10 min to allow phage adsorption. To remove free phages, cells were pelleted by centrifugation at 13,000 × g for 1 min and washed twice with an equal volume of fresh LB medium after removing the supernatant. The infected cultures were aliquoted into multiple tubes and incubated at 37°C with shaking. Samples were collected every 5 min, immediately serially diluted, and spotted onto LB agar overlays (0.5% w/v) mixed with 1% overnight culture of *E. coli* MG1655 supplemented with 10 mM MgCl₂. Data are shown as the mean ± standard deviation (SD) of three biological replicates.

#### Protein expression

For purification, all proteins were expressed in a modified *E. coli* MG1655 strain carrying the DE3 lysogen (Invitrogen, hereafter MG1655-DE3). For Yin1 and Sen3 wild-type and mutants purifications, MG1655-DE3 was co-transformed with two plasmids: a pET-Duet vector encoding the msrRNA, msDNA, and the reverse transcriptase (RT) carrying a HIS-tag in the C-terminus, and a pACYC-Duet vector encoding a N-terminal HA-tagged HEPN and non-tagged HTH proteins, respectively. Cultures were grown at 37 °C in Terrific Broth (TB) medium supplemented with 0.5% glucose, 100 µg/mL ampicillin, and 34 µg/mL chloramphenicol. Cells were grown to an optical density at 600 nm (OD_600_) of 0.6, after which protein expression was induced by the addition of 0.2 mM IPTG for 3 h at 37 °C. An N-terminal His-tagged version of B14T WT or mutants was expressed from a p15a plasmid. Cultures were grown at 37°C in LB medium, supplemented with 0.5% glucose and 34 µg/mL chloramphenicol. Cells grew to an optical density at 600 nm of 0.6 and protein expression was induced by adding 0.15% Arabinose at 18°C, overnight. Upon expression, cells were harvested by centrifugation, flash frozen, and stored at -80°C.

#### Protein purification

Cell pellets were resuspended in lysis buffer containing 20 mM HEPES (pH 7.5), 500 mM NaCl, 20 mM imidazole, 1 mg/mL lysozyme, and one tablet of Complete EDTA-free protease inhibitor cocktail (Roche) per 50 mL. Suspensions were incubated for 40 min at 4 °C with gentle agitation and further lysed by sonication. Lysates were clarified by centrifugation at >10,000 × g for 45 min at 4 °C to remove cellular debris.

The supernatant was loaded onto a 5 mL HisTrap FF Crude column (Cytiva) pre-equilibrated in IMAC-A buffer (20 mM HEPES pH 7.5, 500 mM NaCl, 20 mM imidazole). Bound proteins were eluted using a stepwise imidazole gradient with IMAC-B buffer (20 mM HEPES pH 7.5, 500 mM NaCl, 500 mM imidazole), with the Yin1 complex eluting at approximately 120 mM imidazole. Peak fractions were pooled, concentrated, and further purified by size-exclusion chromatography on a Superose 6 or Superdex 200 Increase 10/300 column (Cytiva) equilibrated in SEC buffer (20 mM HEPES pH 7.5, 150 mM KCl, 0.5 mM TCEP). Fractions containing retron complexes were pooled, concentrated to an A_260_ of 5-10, flash-frozen in liquid nitrogen, and stored at −80 °C. The Sen3 sample was purified following the same procedure, except that the HEPN domain was absent. This preparation was subsequently used for msDNA extraction and sequencing.

B14T was purified using the same procedure, with the inclusion of an additional heparin affinity chromatography step after IMAC to remove contaminating nucleic acids. For this, IMAC fractions containing B14T were pooled, diluted to a final NaCl concentration of 150 mM, and loaded onto a HiTrap Heparin HP column (Cytiva). Proteins were eluted using a linear gradient from 150 mM to 2 M NaCl in 20 mM HEPES pH 7.5 and 0.5 mM TCEP. Peak fractions containing B14T were pooled, concentrated, and subjected to size-exclusion chromatography. In a Superdex 200 increase 10/300 column equilibrated in SEC buffer as described above. Protein identity was confirmed by tryptic digestion followed by mass spectrometry. For B14T, this analysis additionally revealed proteolytic trimming of predicted disordered regions, consistent with purification of the N-terminal nuclease domain (residues 18-295).

#### Extraction of msDNA and library preparation of msDNA

msDNA was isolated from purified Yin1 and Sen3 retron complexes by RNase A treatment followed by phenol-chloroform extraction. Retron-extracted msDNA/msrRNA samples were treated with RNase H (NEB) and DBR1 (OriGene) to remove RNA moieties following the manufacturer’s protocol. Nucleic acids were purified using the Zymo ssDNA/RNA Clean & Concentrator kit and eluted in 10 μL nuclease-free water. Purified ssDNA was subjected to 3′ terminal poly(A) tailing using Terminal Transferase (NEB), and the reaction was terminated by incubation at 70°C for 5 min. After annealing with a Unique Molecular Identifier (UMI)-poly(T9) anchor primer, second-strand synthesis was carried out using Klenow Fragment (3′→5′ exo–) (Thermo Fisher) at 37°C for 30 min, followed by purification using a QIAquick PCR purification kit. Left adapters were annealed and ligated using Blunt/TA Ligase Master Mix (NEB) with a 15 min incubation at room temperature. Indexed libraries were generated using the NEBNext Ultra II DNA Library Prep Kit for Illumina, and PCR amplification was performed for 12 cycles (98°C for 30 s; 12 cycles of 98°C for 10 s and 65°C for 75 s; final extension at 65°C for 5 min). PCR products were purified using AMPure XP beads at a 1.8× bead-to-sample ratio, eluted in 0.1× TE buffer, and stored at −20°C until sequencing (NovaSeq X Plus Series PE150, Novogene).

#### msDNA sequencing data analysis

Adapters were trimmed and reads were quality-filtered with fastp v0.24.0, requiring a minimum read length of 30 nt and a qualified base quality threshold of Q30. Read 1 were used for downstream analyses. PCR duplicates were removed using custom Python scripts by collapsing reads with identical UMIs and retaining one representative read per unique UMI. The deduplicated reads were then mapped to the plasmid reference using Bowtie2 v2.5.4^52^. Mapped reads were retained, sorted, and indexed with SAMtools v1.21^53^ to generate coordinate-sorted BAM files. Per-base coverage across the reference was calculated using BEDTools v 2.31.1^54^ and visualized to identify the msDNA region. To detect cleavage associated read end signal, the 5′ and 3′ ends of primary mapped reads were counted separately at each plasmid position. Fold changes were then calculated for 5′-end and 3′-end counts and visualized across the msDNA region to infer cleavage sites.

#### Electron microscopy and image processing

Yin1 samples were resuspended in imaging buffer (20 mM HEPES pH7.5, 100 mM KCl, 1 mM TCEP). Two sample concentrations were used for imaging, yielding datasets enriched in either ring assemblies or disrupted repeating units (A_260_ = ∼1 and ∼5, respectively). 3 µL were applied to R1.2/1.3 Cu300 mesh grids (Quantifoil) glow-discharged for 60 s at 10 mA (Leica EM ACE200). Grids were plunge-frozen in liquid ethane pre-cooled with liquid nitrogen using a Vitrobot Mark IV (FEI, Thermo Fisher Scientific) under the following conditions: 3 s blotting time, 100% humidity, and 4 °C.

The grids were imaged using EPU v.3.10.0 on a Titan Krios operated at 300keV using a TFS Falcon 4i camera and equipped with a TFS Selectris X energy filter (core facility of integrated bioimaging, CFIB, University of Copenhagen) using counting mode at a magnification of x 165000 (0.728 Å/px) using a total dose of ∼40 e/Å2 using EER format. We used a defocus range spanning −0.6 to −1.8 in 0.2 µm steps. Approximately 3,000 movies were collected for the ring assembly samples (A_260_ ≈ 1; Figure S4A), whereas ∼50,000 movies were collected for the disrupted repeating units (A_260_ ≈ 5; Figure S4B). In both datasets, ∼60% of the movies were acquired with a stage tilt of 40° to enhance map isotropy. All image processing was performed using CryoSPARC v5.0.2^55^. Cryo-EM movies were motion-corrected using patch-based alignment with a 500 Å patch size, and contrast transfer function (CTF) parameters were estimated using patch CTF estimation within CryoSPARC. Micrographs were denoised using the CryoSPARC Micrograph Denoiser^55^ and used for subsequent particle picking.

For the full ring assemblies (A_260_ ≈ 1), particle coordinates were initially determined using the CryoSPARC blob picker (50 Å diameter; Figure S4A). A total of ∼500,000 particles were extracted (box size: 720 pixels, ∼524 Å; downsampled pixel size: 4.1 Å) and subjected to 2D classification, from which ∼173,000 particles were selected. These particles were used to generate an initial model by *ab initio* reconstruction with a single class, following by removal of duplicated particles (minimum distance separation 100 Å) followed by homogeneous refinement. The final 96.927 particles yielded a reconstruction displaying pronounced anisotropy (cFAR = 0.02) with an overall resolution of 7.2 Å, based on the FSC 0.143 criterion. To enable confident assignment of the number of subunits per ring, particles were re-extracted using a larger box size (1000 pixels, ∼729 Å) and subjected to an additional 2D classification.

For the disrupted repeating units (A_260_ ≈ 5; Figure S4B), particles were identified using a circular blob of 100-250Å (CryoSPARC) ^55^. Picking was intentionally performed with high density (minimum interparticle distance of 40 Å) to avoid missing rare particle orientations. Approximately ∼13.4 million particles were extracted using a box size of 720 pixels (∼524 Å) at a down sampled pixel size of 4.1 Å. Following two-dimensional (2D) classification, ∼4 million particles were selected and subjected to ab initio reconstruction using multiple classes. The class exhibiting the highest map quality and most isotropic particle orientation distribution was selected for further processing. Particles from this class were re-extracted at the original pixel size (0.728 Å). To improve Fourier sampling, additional rounds of 2D classification, three-dimensional (3D) classification without alignment, and particle orientation balancing were carried out. This yielded a final dataset of 525.790 particles, for which the defocus values were further refined using local motion correction and local and global CTF refinement. The selected particles were then subjected to non-uniform refinement with Ewald sphere curvature correction, resulting in an improved and more isotropic reconstruction (cFAR = 0.28) with an overall resolution of 3.24 Å (FSC criterion 0.143).

Final maps were sharpened using either Phenix^56^ AutoSharpen (*B* factor: -138.72 Å^2^) or DeepEMhancer^57^ to facilitate map interpretation and model building. Unsharpened primary maps, and half maps were deposited on the EMDB and their local resolution was calculated in CryoSPARC using a a FSC criterion 0.5 (Figure S5C and S5I)

#### Structure modelling

AlphaFold3 (AF3) models^28^ comprising the HEPN, KH-HTH, and RT proteins of the Retron-Yin1 were rigid-body fitted into the cryo-EM maps of the disrupted repeating units using ChimeraX^58^ (Figure S3F and S6D). The resulting model was manually inspected and adjusted in ISOLDE^59^ and Coot^60^. The msrRNA and msDNA components were built using a combination of AF3-derived and idealized nucleic acid models generated in Coot^59^. The complete model containing all proteins and nucleic acids was further refined in ISOLDE^59^ in Phenix^56^. The model of the refined repeating unit was used to generate a model of the full nine-subunit ring by fitting it into two consecutive segments of the 9-mer cryo-EM map (EMDB-58248) and propagating the symmetry ninefold in ChimeraX^58^. All molecular figures were generated using ChimeraX^58^.

#### Structural comparisons and evolutionary conservation analysis

Structural homologs of Yin1 HEPN and B14T were identified using Foldseek^61^. Searches for Yin1 HEPN domain homologs identified HEPN–MNT systems as the closest matches. B14T homologs included AdnA and RecB nucleases. Homologous domains were subsequently superposed in ChimeraX^58^ to calculate root mean square deviation (RMSD) values. Evolutionary conservation scores for msDNA were calculated using ConSurf^62^, based on the multiple sequence alignment used to generate the consensus secondary structure of the type IX ncRNA. Conservation scores were subsequently mapped onto the msDNA cartoon.

#### Phage DNA extraction

Phage genomic DNA was extracted as previously described. Briefly, filter-sterilized phage lysates were treated with DNase I (QIAGEN) and RNase A (QIAGEN) at 37°C for 1.5 h to remove contaminating bacterial nucleic acids. EDTA was then added to a final concentration of 20 mM. Phage capsids were digested by incubation with Proteinase K (QIAGEN) at 56°C for 1.5 h. DNA was then purified using the DNeasy Blood & Tissue Kit (Qiagen) according to the manufacturer’s instructions. In brief, lysates were mixed with Buffer AL, incubated at 70°C, followed by ethanol addition and loading onto DNeasy Mini spin columns. Columns were washed sequentially with Buffers AW1 and AW2, and DNA was eluted in AE buffer^63^.

#### Phage escaper analysis

Phage genomes were sequenced to identify escaper mutations associated with the escape phenotype. Genomic DNA libraries were prepared using the Nextera XT DNA Library Preparation Kit (Illumina) according to the manufacturer’s instructions. Paired-end sequencing (2 × 300 bp) was performed in-house on an Illumina MiSeq platform using the MiSeq Reagent Kit v3. Raw reads were adapter-trimmed and quality-filtered using fastp v0.24.0^64^. High-quality reads were assembled de novo with Unicycler^65^. Genome annotation was performed using Pharokka^66^. To identify genetic variants, trimmed reads were mapped to the corresponding reference phage genome and mutations were called using Breseq^67^. Detected mutations were further examined to determine affected genes and potential functional consequences.

#### Sequence analysis

B14T homologs were identified using hmmsearch in HMMER v3.4^68^ with the DUF2800 Pfam HMM profile PF10926 as the query against the INPHARED^69^ database of RefSeq complete phage genomes (release 1Mar2024), filtered with an E-value of 1e-3. To construct the B14T phylogenetic tree, sequences longer than 150 amino acids were retained, and redundant sequences were removed using CD-HIT v4.8.1^70^ with a sequence identity threshold of 100% and a length difference cutoff of 100%. The representative sequences were aligned using MAFFT v7.525^71^ with the FFT-NS-1000 strategy, and the alignment was trimmed using trimAl v1.5.0^72^ by removing columns containing more than 30% gaps. Phylogenetic tree construction was performed using FastTree v2.1.1^73^ with gamma-distributed site rates and the WAG evolutionary model. The tree was visualized using the Python package ETE Toolkit.^74^ The sequence logo of the region corresponding to the B14T PD-(D/E)XK motif was generated from the trimmed alignment of B14T homologs using WebLogo v3.7.12^75^ with the chemistry colour scheme.

Retron IX homologs were retrieved from NCBI refseq non-redundant protein database, and the corresponding covariance model was constructed as described previously^12^. Retron IX components and their respective homologs were aligned using MAFFT v7.525^71^ with the FFT-NS-1000 strategy, and the resulting sequence alignments were visualized with ESPript^76^. The phylogenetic tree of retron IX reverse transcriptases was reconstructed using the sequence alignment and tree-building workflow described above. To determine whether a retron IX system was associated with a defence island, known defence systems within 20,000 bp upstream and downstream of the RT were identified using PADLOC v2.0.0^77^, excluding the retron IX system itself. To assess whether retron IX systems were associated with prophages or plasmids, the genetic elements were detected using the geNomad v1.8.0^78^ end-to-end pipeline within 75,000 bp upstream and downstream of the RT.

#### Toxicity assays with B14T

To analyse the toxicity triggered by the coexpression of retron IX and B14T, single colonies of *E. coli* MG1655 harbouring Sen3 and empty vector, B14T wild-type or catalytic mutant (D177A/K179A) were grown at 37°C overnight in LB supplemented with 2 µg/mL spectinomycin, and 25 µg/mL chloramphenicol and 0.3% glucose. Overnight cultures were diluted 1:100 into the same medium and grown to an OD₆₀₀ of 0.2. Cells were then washed with clean LB and resuspended to the same volume. For induction, 100 µL of the resuspended culture was mixed with 100 µL of induction medium preloaded in 96-well plates, yielding final concentrations of 0.15% arabinose, 2 µg/mL spectinomycin, and 25 µg/mL chloramphenicol. The plate was incubated at 37°C with shaking at 200 rpm in a BioTek Synergy H1 microplate reader, and OD₆₀₀ was measured every 5 min.

#### B14T-mediated msDNA cleavage assay *in vitro*

B14T-mediated Yin1 msDNA cleavage reactions were performed by incubating 1 µM Yin1 with 1.7 µM B14T in cleavage buffer (20 mM HEPES pH 7.0, 100 mM KCl, 5 mM MgCl₂, 0.5 mM TCEP) at 37 °C. Samples were collected at the indicated time points, and reactions were quenched by addition of urea loading buffer (final concentration 10 mM Tris pH7.5, 4M Urea, 5 mM EDTA) followed by denaturation at 95°C for 10 min. Cleavage products were analyzed on 15% TBE-urea gels stained with GelRed (Millipore) or SYBR Safe (Invitrogen^TM^)

#### Pull down assays

For pull down experiments with Yin1 and Yin1/Sen3 incubated with B14T, 50 mL of cultures were grown at 37°C in a liquid LB media and expressed as indicated above (protein purification). Cells were harvested by centrifugation, flash frozen, and stored at -80°C. Bacterial pellets were resuspended in 1mL pull down buffer (20 mM HEPES, pH 7.5, 300 mM KCl, 5 mM MgCl₂, and 0.1% Triton X-100). Cells were disrupted by sonication, and lysates were clarified by centrifugation at 4°C for 10 min at x14.000 g to separate the soluble fraction from the insoluble fraction and cell debris. B14T and retron lysates were mixed at 1:1(v/v) ratio and incubated 30 min at 25°C with gentle agitation. Following incubation, samples were centrifuged at 4°C for 1 min at maximum speed to remove aggregates, and the clarified supernatant was transferred to 20 µL Ni-NTA agarose beads (Qiagen) pre-equilibrated in pull down buffer. After incubation with rotation for 10 min at 4°C, beads were washed four times with 500 µL pull down buffer. After the final wash, beads were resuspended in approximately 60 µL pull down buffer and split for downstream protein (SDS-PAGE electrophoresis and Western Blot) and nucleic acid (TBE-Urea electrophoresis) analysis.

#### Electrophoresis and Western blot

Protein samples were denatured in SDS-PAGE loading buffer at 95°C for 5 min, prior to separation on NuPAGE^TM^ 4–12% Bis-Tris gels (Invitrogen^TM^). Proteins were transferred to nitrocellulose membranes using iBlot™ Transfer Stacks (Invitrogen^TM^). Membranes were blocked with PBST containing 5% skim milk powder and incubated with HRP-conjugated primary anti-Penta-His (Qiagen) or anti-HA (Roche) (1:1000 v/v diluti. After washing with PBST, chemiluminescent signal was detected using SuperSignal™ West Pico PLUS Chemiluminescent Substrate (Thermo Scientific^TM^) or SuperSignal^TM^ West Atto Ultimate Sensitivity Chemiluminescent Substrate (Thermo Scientific^TM^). Samples for nucleic acid analysis were mixed with urea loading dye (final concentration 10 mM Tris pH7.5, 4M Urea, 5 mM EDTA) followed by denaturation at 95°C for 10 min. When specified (“+RNAse A”) samples were treated with 100 μg/mL of RNAse A at 20°C for 10 min. Nucleic acid sizes were estimated by comparison to an ss20 DNA ladder, ss50 DNA ladder (Simplex Sciences) or a homemade ladder containing 50-90 nt bands with intervals of 10 nt. Denaturing polyacrylamide gel electrophoresis was performed using 15% TBE-Urea Gels (Thermo Scientific^TM^), followed by staining with GelRed (Millipore) or SYBR Safe (Invitrogen^TM^) for detection.

#### Quantification of DNA and protein electrophoresis

In pull-down assays, msDNA abundance was quantified by densitometry of the uncleaved (upper) band. Signals were first corrected for RT abundance, determined by densitometry of western blots from the same samples, and then scaled relative to the most intense uncleaved band (−B14T; set to 1). Finally, values were normalized internally across retron types (Yin1 and Yin1/Sen3) to account for differences between −B14T and +B14T conditions. HEPN abundance was quantified in the same manner by densitometry of western blot bands.

#### RNA extraction and sequencing

*E. coli* MG1655 harbouring empty vector control, active, or inactive constructs for Sen3 and Bas14_0045 (Sen3_Mut, HEPN H64A; B14T_Mut, D177A/K179A) were grown at 37°C overnight in LB supplemented with 2 µg/mL spectinomycin, and 25 µg/mL chloramphenicol and 0.3% glucose. Overnight cultures were diluted 1:100 into the same medium and grown to an OD₆₀₀ of 0.2. Cells were then washed with clean LB and resuspended to the same volume with LB supplemented with 2 µg/mL spectinomycin, and 25 µg/mL chloramphenicol and 0.15% arabinose. The culture was then incubated at 37°C with shaking for 60 min. Total RNA was extracted using the Zymo Research Quick-RNA Fungal/Bacterial Miniprep Kit with on-column DNase I treatment to retain RNAs >17 nt and remove residual msDNA. Small RNAs (17–200 nt) were enriched using the Zymo Research RNA Clean & Concentrator-5 Kit. To prepare RNA termini for downstream adapter ligation and library construction, enriched small RNAs were treated with 40 U of T4 Polynucleotide Kinase (PNK, NEB) for 30 min at 37°C and 20 U of RNA 5′ Pyrophosphohydrolase (RppH, NEB) for 30 min at 37°C prior to library construction. Each treatment step was followed by cleanup using a Zymo RNA Clean & Concentrator-5 kit according to the manufacturer’s instructions. Libraries were prepared using the NEBNext Multiplex Small RNA Library Prep Set for Illumina (E7300) according to the manufacturer’s instructions and sequenced using paired-end 150 bp reads (PE150) on an Illumina platform (Novogene).

#### RNA sequencing data analysis

Raw reads were trimmed using fastp v0.24.0 (--detect_adapter_for_pe, --disable_quality_filtering, --n_base_limit 0, and --length_required 17). Reads were then aligned to *E. coli* K-12 MG1655 reference genome (U00096.3) using Bowtie2 v2.5.4. Mapped reads were converted to BAM format, filtered with SAMtools v1.21 to remove unmapped, secondary, and supplementary alignments (-F 2308), and then sorted and indexed. Read 1 were used for downstream analysis. Read end coordinates were extracted from sorted BAM files using BEDTools v2.31.1. Internal read end enrichment analysis was performed by mapping read end coordinates to annotated genomic features in *E. coli* K-12 MG1655 (U00096.3). For tRNA anticodon-loop cleavage analysis, tRNA sequences were extracted from the U00096.3 reference annotation and anticodon positions were located by tRNAscan-SE v2.0.12^79^. Read end signals were normalized to count per million mapped reads (CPM). Cleavage signals were calculated as the log_2_ fold change in read end counts in Sen3-B14T relative to Sen3_Mut-B14T. To avoid undefined values during fold change calculation, a pseudocount of 1 was added to read end counts at all genomic positions. Mean positional log_2_ fold change values were then mapped onto a schematic representation of the tRNA anticodon loop. To better visualize cleavage sites, the colour scale was set to start from 0. To identify cleavage target, we adapated Python scripts as previously described^30^. Briefly, read end counts fold changes were calculated at each genomic position by comparing the active sample (Sen3-B14T) against all control samples. Genomic positions with read end counts fold change greater than 30 in Sen3-B14T relative to the controls were identified as candidate cleavage sites. The corresponding positions were then annotated using the reference genome annotation. To assess whether the detected targets exhibited sequence preference, the anticodon sequence together with seven nucleotides upstream and downstream of were extracted as anticodon loop sequence from the detected tRNAs and made sequence logo compares to those of all E. coli tRNA repertoire. To visualize the overall fold change signal, positional fold change values of Sen3-B14T relative to Sen3_Mut-B14T read end counts were mapped onto the schematic secondary structure of tRNA-Gly(UCC).

#### tRNA cleavage assay *in vitro*

Yin1 wild-type and the R116A HEPN mutant were incubated at 1 µM with either 2 µM B14T WT or the inactive B14T mutant (D177A/K179A) in buffer containing 20 mM HEPES (pH 7.5), 50 mM KCl, 2 mM MgCl₂, and 1 mM TCEP for 30 min at 37 °C. Following msDNA cleavage, 0.4 µM N-terminal FAM-labelled tRNA-Gly (UCC; IDT) was added, and reactions were continued at 30 °C. Aliquots were collected at defined time points (0, 10, and 30 min), and reactions were quenched by addition of urea loading dye (final concentrations: 10 mM Tris-HCl pH 7.5, 4 M urea, 5 mM EDTA), followed by denaturation at 95 °C for 10 min. Samples were resolved on 15% TBE-urea polyacrylamide gels and analysed by FAM fluorescence imaging. Subsequently, gels were stained with GelRed for 10 min to visualize msrRNA and msDNA bands.

