## Supplementary Figures for "A DNA-scaffolded Retron Ring Mediates Antiphage Immunity"

Figure S1

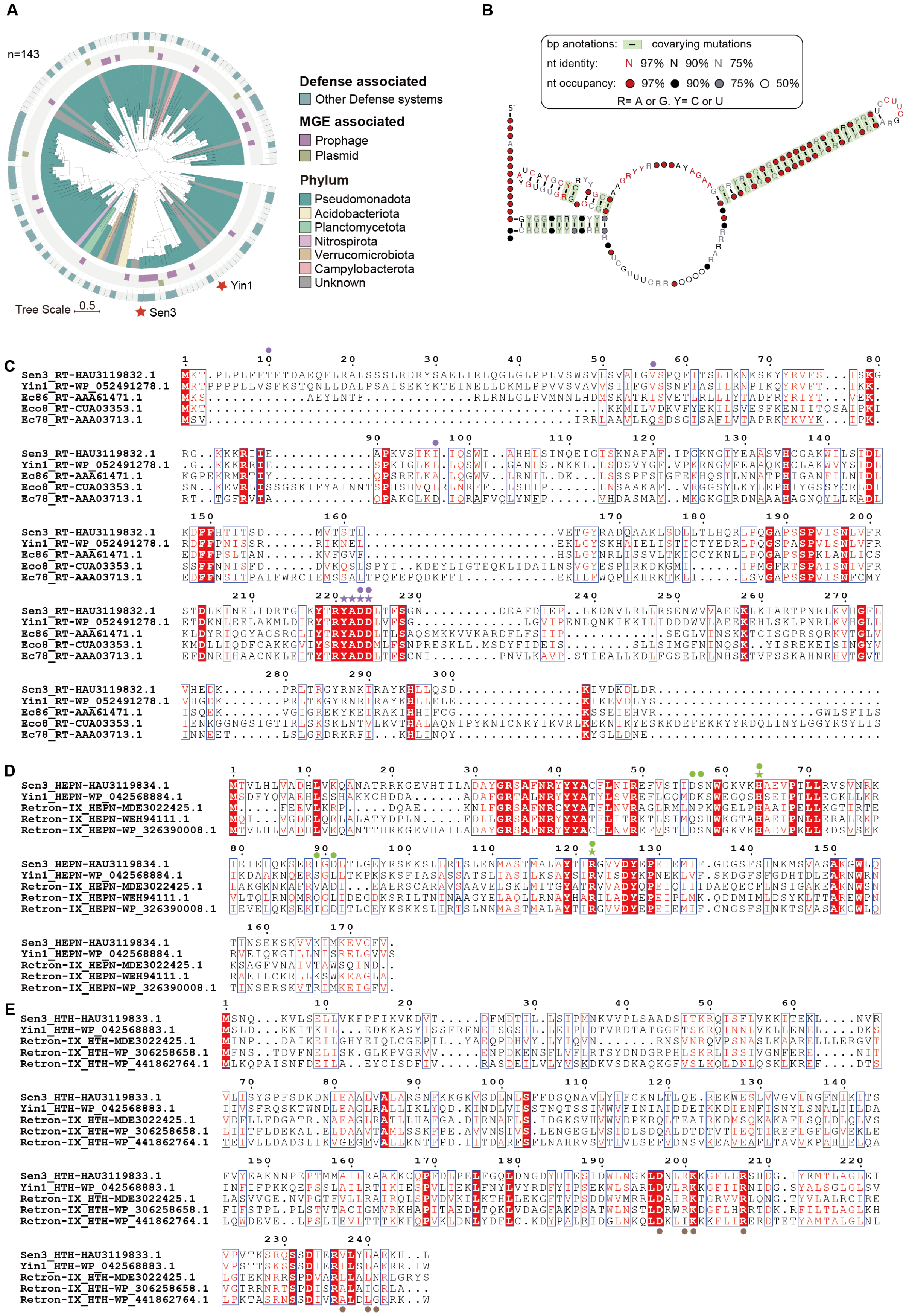
A
n=143
Defense associated
Other Defense systems
MGE associated
Prophage
Plasmid
Phylum
Pseudomonadota
Acidobacteriota
Planctomycetota
Nitrospirota
Verrucomicrobiota
Campylobacterota
Unknown
Yin1
Sen3
Tree Scale 0.5
B
bp anotations: covarying mutations
nt identity: N 97% N 90% N 75%
nt occupancy: 97% 90% 75% 50%
R= A or G. Y= C or U
C
Sen3_RT-HAU3119832.1
Yin1_RT-WP_052491278.1
Ec86_RT-AAA61471.1
Eco8_RT-CUA03353.1
Ec78_RT-AAA03713.1
D
Sen3_HEPN-HAU3119834.1
Yin1_HEPN-WP_042568884.1
Retron-IX_HEPN-MDE3022425.1
Retron-IX_HEPN-WEH94111.1
Retron-IX_HEPN-WP_326390008.1
E
Sen3_HTH-HAU3119833.1
Yin1_HTH-WP_042568883.1
Retron-IX_HTH-MDE3022425.1
Retron-IX_HTH-WP_306258658.1
Retron-IX_HTH-WP_441862764.1

**Figure S1.- Distribution of IX retrons, ncRNA secondary structure, and multiple sequence alignments**

**A)** Phylogenetic tree of type IX retron reverse transcriptases (RTs). Leaf colours indicate bacterial phylum. The three outer rings, from outer to inner, indicate association with known defence systems, prophages, and plasmids, respectively. Red stars indicate the type IX retron RTs characterized in this study. **B)** Consensus ncRNA secondary structure of Sen3. **(c-e)** Sequence alignments of the RT, HEPN and KH-HTH domains of Sen3, Yin1 and representative homologues, respectively. Residues marked with a star correspond to catalytic residues in the RT and HEPN proteins, while circles indicate residues mutated in this study for functional analysis.

Figure S2

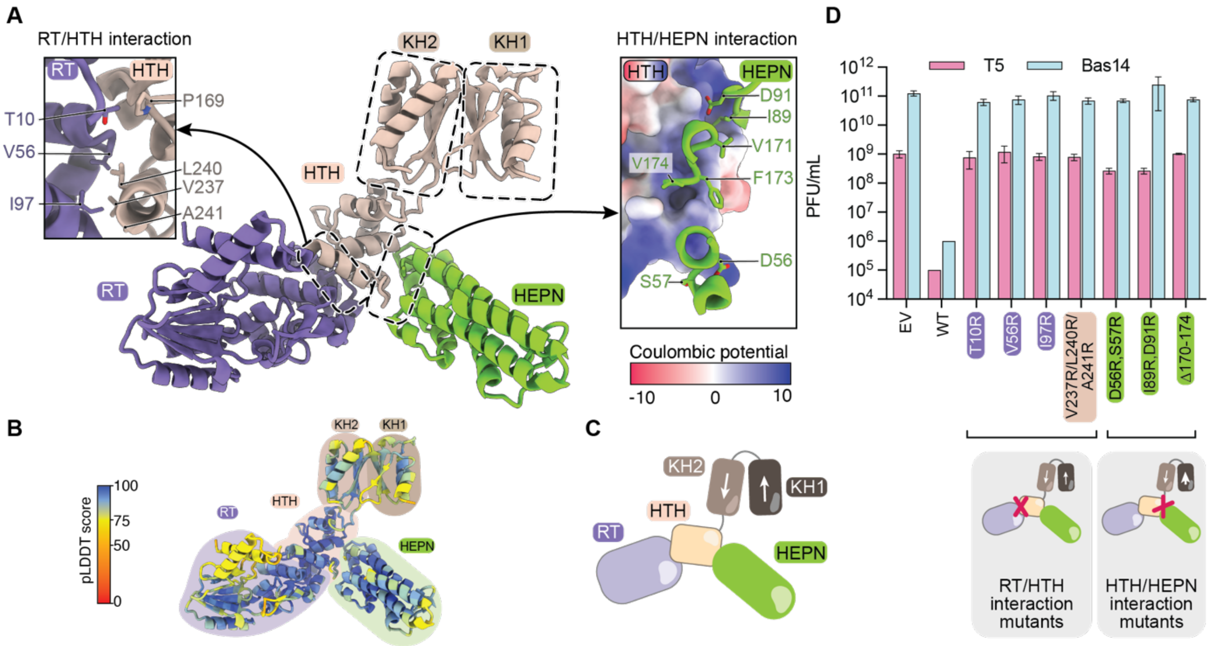


**Figure S2.- Structural model of retron IX components and functional validation.**
**A)** AlphaFold3 (AF3) model of the RT, KH1-KH2-HTH, and HEPN proteins (ipTM: 0.74, pTM: 0.73). Insets show close-up views of the interaction interfaces between the three modules. The electrostatic surface potential of the HTH domain is plotted on the surface on the right. **B)** Predicted Local Distance Difference Test (pLDDT) confidence scores corresponding to the AF3 model shown in (A). **C)** Cartoon representation of the AF3 model in (A). **D)** Plaque-forming units (PFUs) of T5 and Bas14 phages infecting *E. coli* MG1655 strains carrying an empty vector or expressing Sen3 wild-type and variants harbouring mutations at the RT-HTH and HTH-HEPN interfaces. Data represent mean ± SD from three biological replicates.

Figure S3

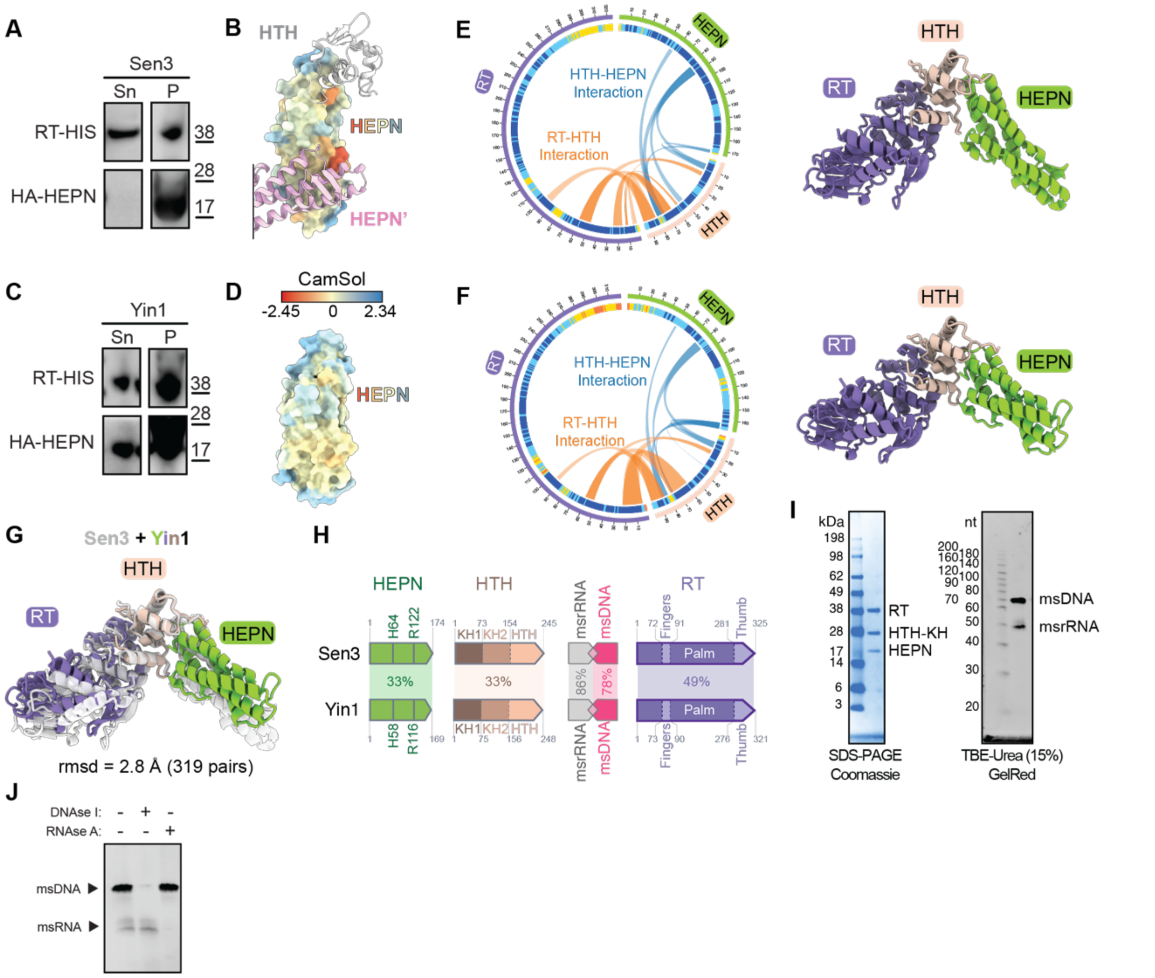


**Figure S3.- Structural and biochemical comparison of Sen3 and Yin1 retrons.**
**A)** Expression and solubility analysis of Sen3 RT and HEPN (H64A) proteins. The Sen3 HEPN domain is largely absent from the soluble fraction following cell lysis, as detected by western blot using an anti-HA antibody. **B)** Structurally corrected CamSol scores mapped onto the surface of the Sen3 HEPN domain. The HTH domain and the second HEPN protomer required to build the catalytically competent dimer are also shown, based on AF3 models. **C)** Expression and solubility analysis of Yin1 RT and HEPN (R116A) proteins. The Yin1 HEPN domain is present in the soluble fraction following cell lysis, as detected by western blot using an anti-HA antibody. **D)** Structurally corrected CamSol scores mapped onto the surface of the Yin1 HEPN domain. (E, F) AF3 models of Sen3 (E) (ipTM: 0.77, pTM: 0.79) and Yin1 (F) (ipTM: 0.73, pTM: 0.73) RT-HTH-HEPN assembly (right). Interaction contacts between RT, HTH, and HEPN domains are shown as a wheel diagram (left) based on the AF3 models obtained with AlphaBridge[51] (left). **G)** Structural superposition of Sen3 and Yin1 AlphaFold models (RMSD: 2.80 Å across 319 pairs). **H)** Comparison of Sen3 and Yin1 retrons. Domain boundaries and catalytic residues are indicated. Amino acid and nucleotides identities between different components are shown as percentages. **I)** Purification of the Retron-Yin1, showing RT, HTH, and HEPN proteins together with msrRNA and msDNA. **J)** TBE-urea gel analysis of msrRNA and msDNA before and after treatment with DNase I and RNase A. The gel images in (A) and (C) are representative of more than three biological replicates. The gel images in (I) and (J) are representative of more than three independent purifications.

Figure S4

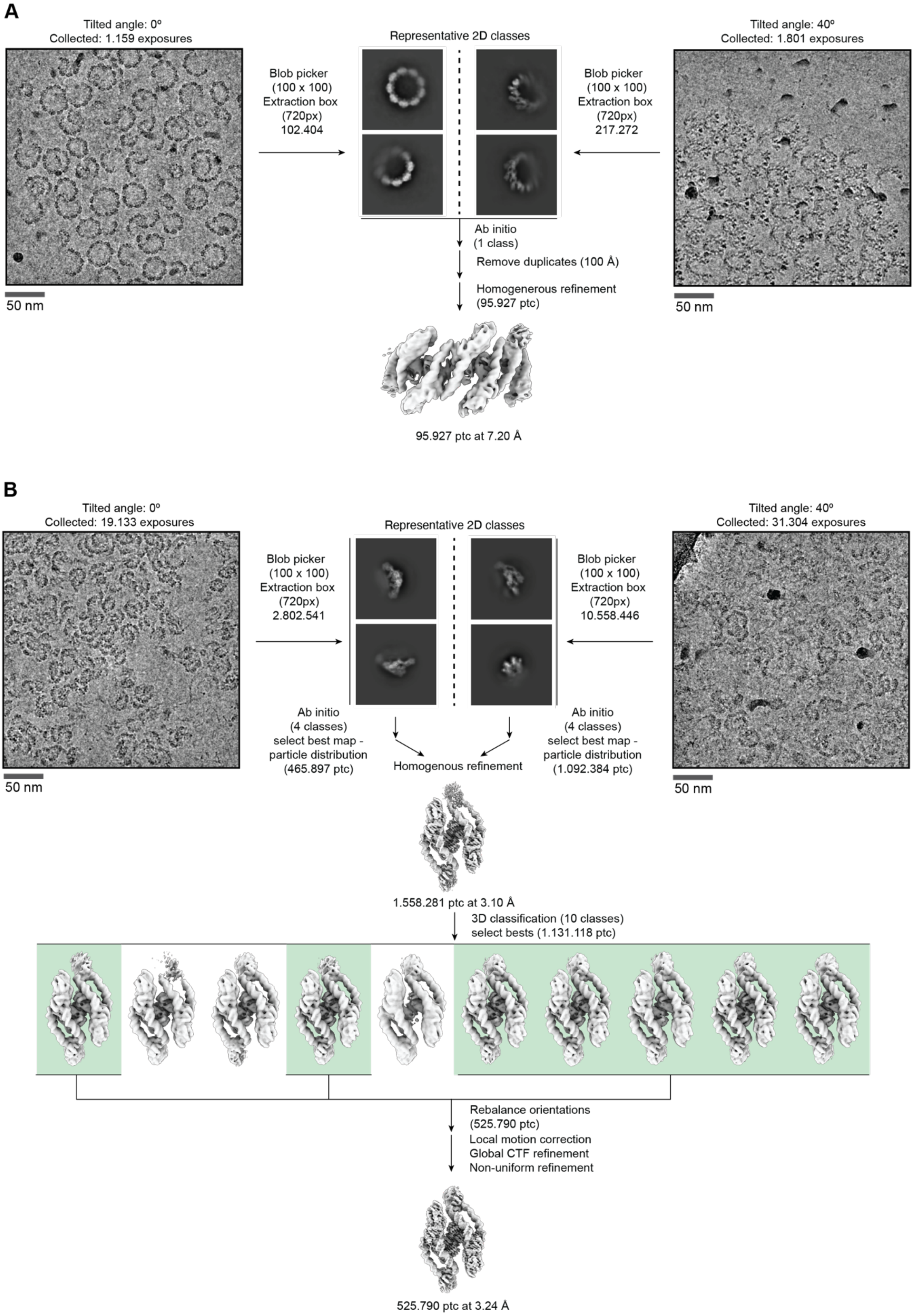


**Figure S4.- Cryo-EM workflows.**
**A)** CryoEM processing workflow of the full rings. **B)** CryoEM processing workflow of the disassembled particles.

Figure S5

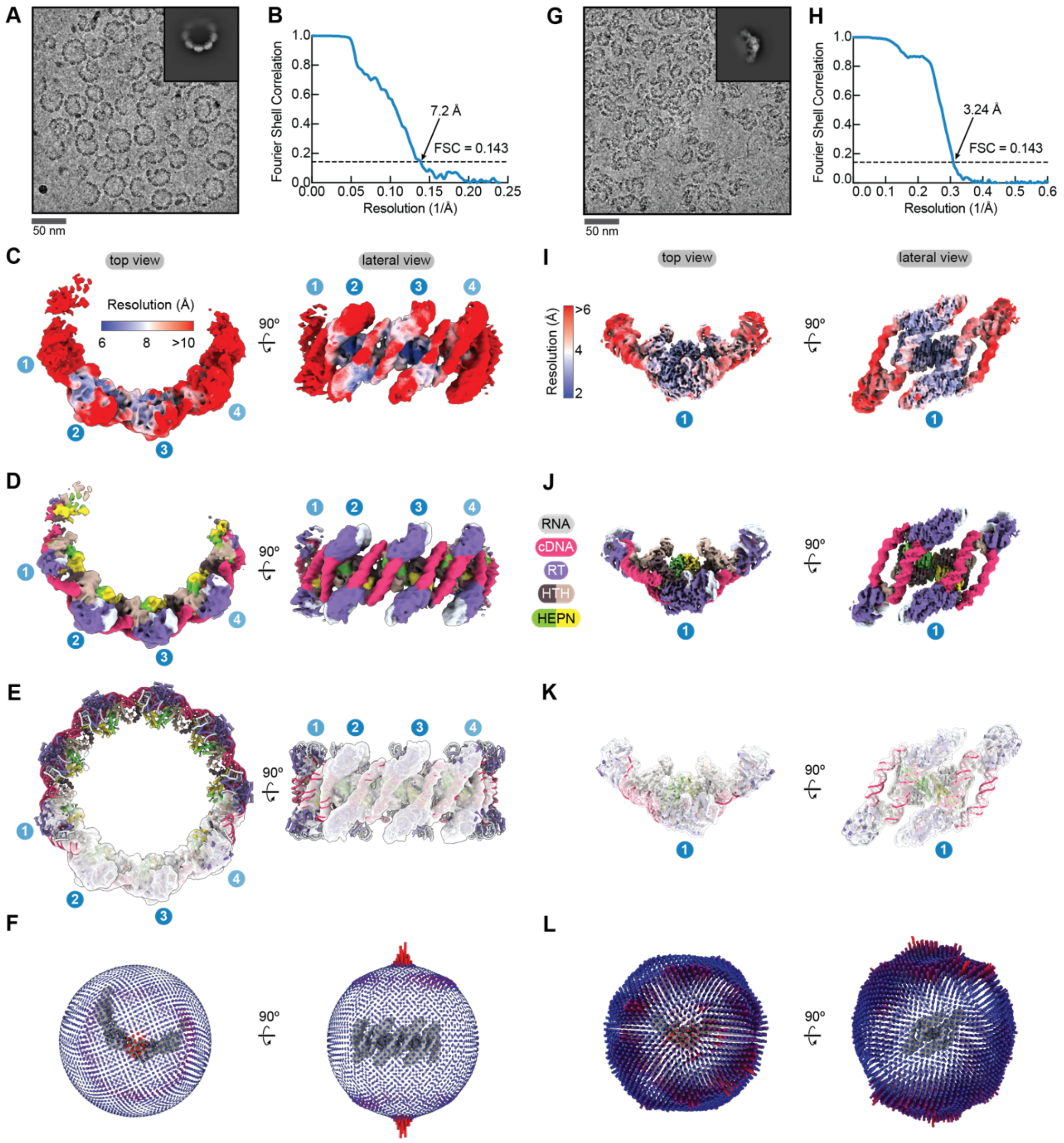


**Figure S5.- Cryo-EM maps of Retron-Yin1 samples.**

**A)** Representative micrograph of Yin1 at low concentration, showing the assembly of ring-like structures. A representative 2D class average of the top view of the complex is shown in the inset. **B)** Fourier shell correlation (FSC) analysis estimating the resolution of the cryo-EM map obtained under low protein concentration conditions, as shown in (A). **C)** Cryo-EM map of the retron ring at low concentration, coloured by local resolution (FSC criteria 0.5). **D)** Cryo-EM map of the retron ring coloured according to the different components. **E)** Fit of the 9-mer retron ring model into the cryo-EM map, shown as a transparent surface. **F)** Angular distribution of retron ring particles contributing to the final reconstruction. The map and angular distribution are shown in the same orientation. **G)** Representative micrograph of Yin1 at high concentration, showing the disassembled repeating units. A representative 2D class average of the top view of the complex is shown in the inset. **H)** FSC analysis estimating the resolution of the cryo-EM map obtained under high protein concentration conditions, as shown in (F). **I)** Cryo-EM map of the retron disassembled repeating units at low concentration, coloured by local resolution. **J)** Cryo-EM map of the retron disassembled repeating units at low concentration, coloured according to the different components. **K)** Fit of the disassembled repeating unit model into the cryo-EM map, shown as a transparent surface. **L)** Particle angular distribution of the disassembled retron ring segments contributing to the final reconstruction. The map and angular distribution are shown in the same orientation.

Figure S6

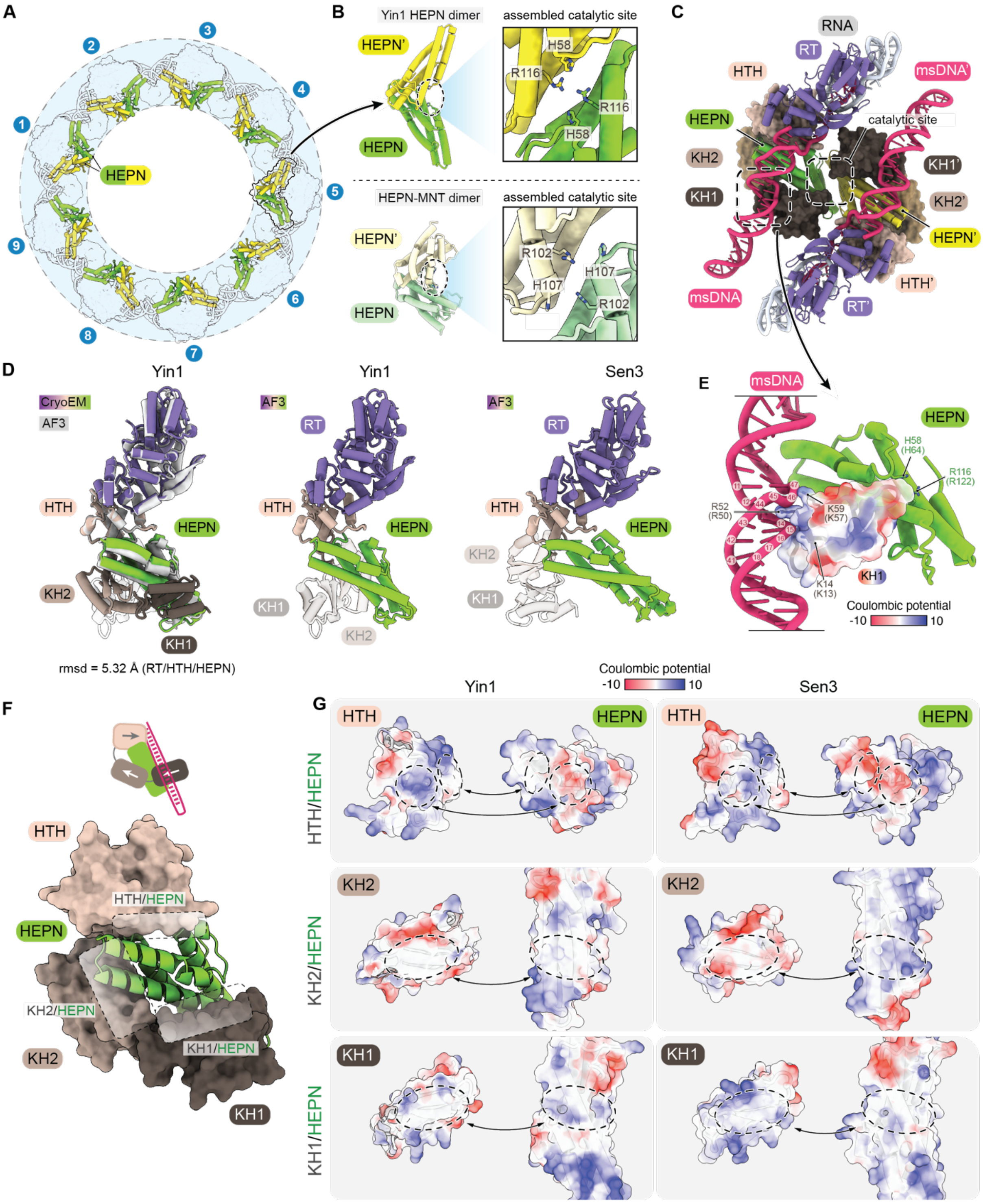


**Figure S6.- Structural details of Yin1.**

**A)** Top view of the Yin1 ring showing HEPN dimers in the inner layer, coloured yellow and green, with the remaining components shown transparently. **B)** Comparison of Yin1 and HEPN-MNT toxin-antitoxin system[29] (PDB:7AE9) HEPN proteins. Insets highlight the assembly of the composite catalytic sites, with strictly conserved catalytic residues shown as sticks. **C)** Molecular representation of the Yin1 repeating unit, composed of a dimer of the HEPN domain, with each protomer bound to one copy of HTH, RT, msrRNA, and msDNA. **D)** Left: Structural superposition of the experimental model (coloured) and the AlphaFold3 (AF3) model of RT, KH1–KH2–HTH, and HEPN domains (RMSD = 5.32 Å; calculated over the RT,

HTH, and HEPN domains). Middle: AF3 model of Yin1. Right: AF3 model of Sen3. **E)** Detailed view of the interactions between msDNA and the KH1 domain, highlighting three conserved basic residues shown as sticks. The electrostatic potential of the KH1 domain is displayed on the protein surface with transparency. **F)** Close-up view of the interactions between the HTH, KH1, and KH2 domains with the HEPN protein **G)** Comparison of the molecular interfaces formed by HTH, KH1, and KH2 domains with the HEPN protein in the Yin1 structure and Sen3 AF3 predicted models. AF3 models of Sen3 domains were superposed onto the Yin1 experimental structure for visualization. Protein surfaces are coloured by electrostatic potential.

Figure S7

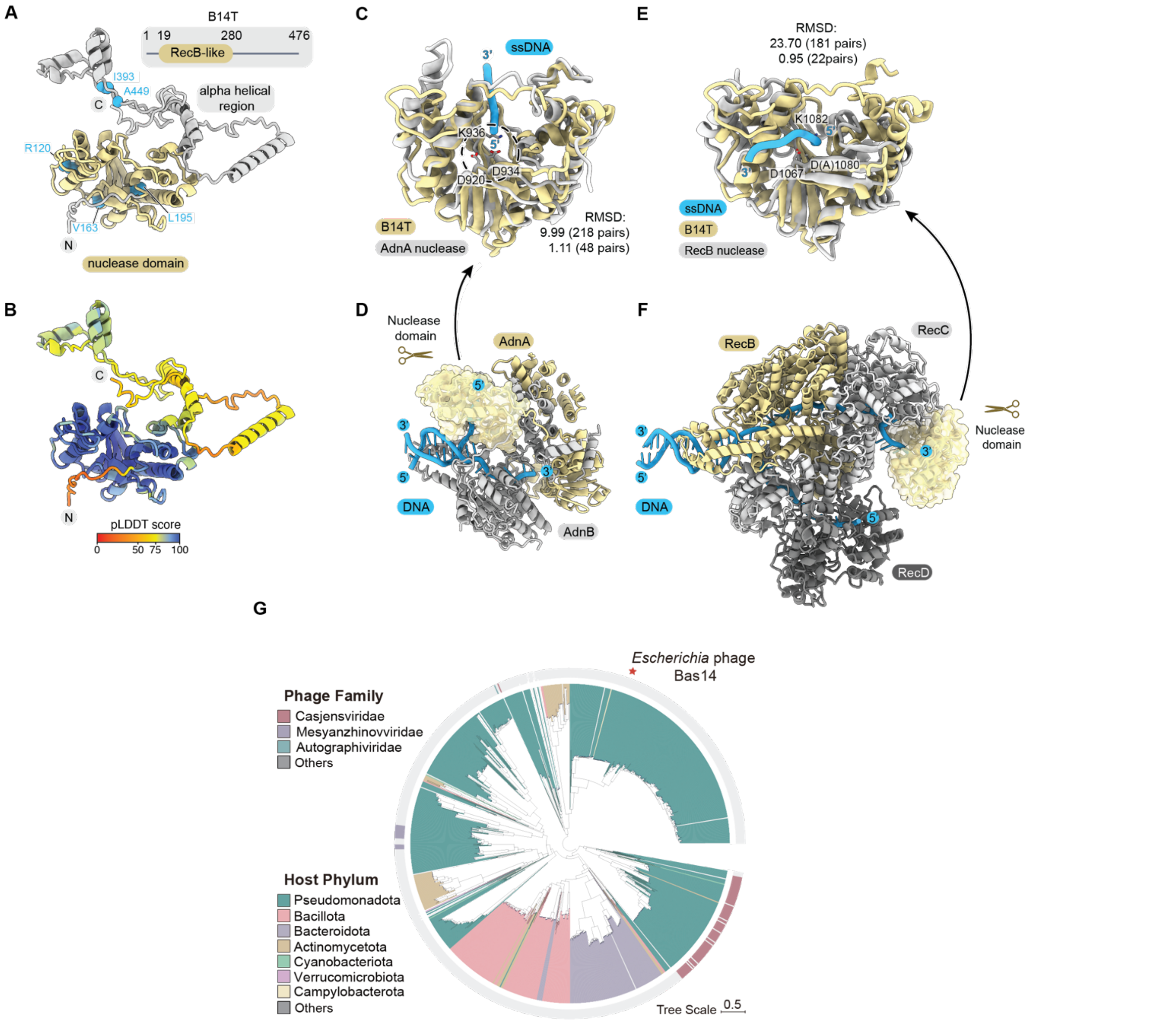


**Figure S7.- Structural comparisons between B14T and homolog nucleases.**
**A)** AlphaFold3 (AF3) model of the B14T protein. The nuclease domain (residues 10–280) is shown in gold, and residues affected by the phage resistant mutations are shown as blue spheres. **B)** Predicted Local Distance Difference Test (pLDDT) scores mapped onto the B14T AF3 model. **C)** Structural superposition of B14T and the AdnA nuclease (PDB:6PPU; RMSD: 9.99 Å across 218 pairs; 1.11 A across 48 pairs). **D)** Molecular model of the AdnA/B complex bound to DNA (PDB:6PPU). The nuclease domain of AdnA is highlighted as a transparent surface. **E)** Structural superposition of B14T and the RecB nuclease (PDB: 6SJB; RMSD: 32.70Å across 181 pairs; 0.95 Å across 22 pairs). **F)** Molecular model of the RecBCD complex bound to DNA (PDB: 6SJB). The nuclease domain of RecB is highlighted as a transparent surface. **G)** Phylogenetic analysis of phage DUF2800 containing nuclease retrieved from the RefSeq complete phage genomes database (INPHARED).

Figure S8

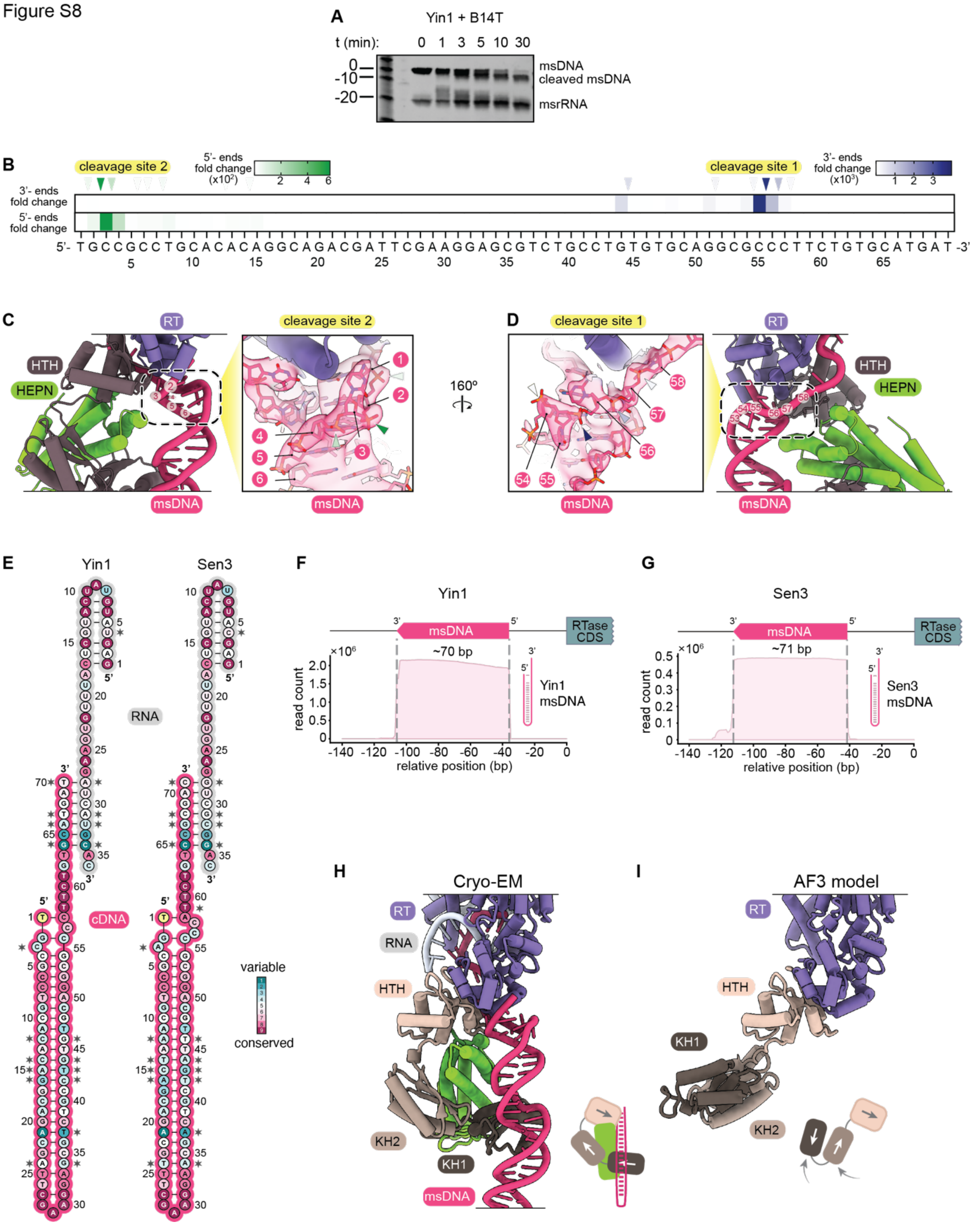


**Figure S8.- Analysis of Yin1 cleavage by B14T.**

**A)** Time-course cleavage assay of Yin1 msDNA by the B14T nuclease. Samples were collected at indicated time points and analyzed by TBE-urea gel electrophoresis. The gel is representative of three independent experiments. **B)** Fold change of 5′ and 3′ cleavage events mapped onto the Yin1 msDNA sequence, as determined by next-generation sequencing. Cleavage sites 1 and 2 are indicated. **C)** Cleavage site 2 shown in the cryo-EM-derived model (left) and in a close-up view (right). **D)** Cleavage site 1 shown in the cryo-EM-derived model (right) and in a close-up view (left). **E)** Schematic representation of Yin1 and Sen3 msrRNA and msDNA, with bases coloured according to evolutionary conservation. Nucleotides that differ between the two retrons are marked with an asterisk. **F-G)** Read counts and coverage of Yin1 (F) and Sen3 (G) msDNAs. The length of the most abundant msDNA species is indicated above each plot. **H)** Molecular model of a single unit comprising 1x RT, msrRNA, HTH, and HEPN, based on the Cryo-EM Yin1 map. **(I)** AlphaFold3 model of the RT-HTH-KH1-KH2 interaction, showing an alternative conformation of the KH1-KH2 tandem.

Figure S9

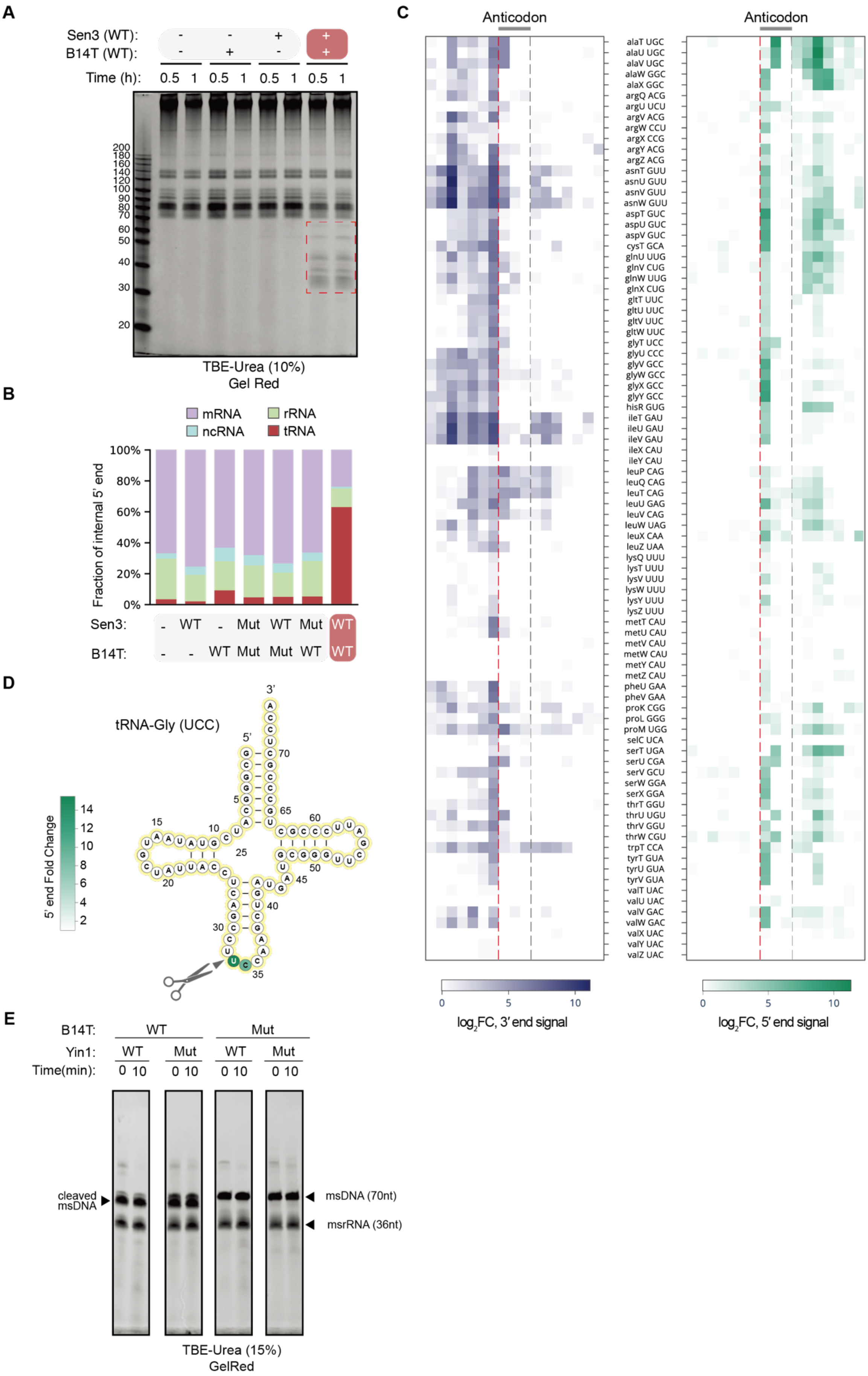
A
Sen3 (WT): - - + +
B14T (WT): - + - +
Time (h): 0.5 1 0.5 1 0.5 1 0.5 1
200
180
160
140
120
100
90
80
70
60
50
40
30
20
TBE-Urea (10%)
Gel Red
B
mRNA
rRNA
ncRNA
tRNA
Fraction of internal 5' end
100%
80%
60%
40%
20%
0%
Sen3: - WT - Mut WT Mut WT
B14T: - - WT Mut Mut WT WT
C
Anticodon
Anticodon
alaT UGC
alaU UGC
alaV UGC
alaW GGC
alaX GGC
argQ ACG
argU UCU
argV ACG
argW CCU
argX CCG
argY ACG
argZ ACG
asnT GUU
asnU GUU
asnV GUU
asnW GUU
aspT GUC
aspU GUC
aspV GUC
cysT GCA
glnU UUG
glnV CUG
glnW UUG
glnX CUG
gltT UUC
gltU UUC
gltV UUC
gltW UUC
glyT UCC
glyU CCC
glyV GCC
glyW GCC
glyX GCC
glyY GCC
hisR GUG
ileT GAU
ileU GAU
ileV GAU
ileX CAU
ileY CAU
leuP CAG
leuQ CAG
leuT CAG
leuU GAG
leuV CAG
leuW UAG
leuX CAA
leuZ UAA
lysQ UUU
lysT UUU
lysV UUU
lysW UUU
lysY UUU
lysZ UUU
metT CAU
metU CAU
metV CAU
metW CAU
metY CAU
metZ CAU
pheU GAA
pheV GAA
proK CGG
proL GGG
proM UGG
selC UCA
serT UGA
serU CGA
serV GCU
serW GGA
serX GGA
thrT GGU
thrU UGU
thrV GGU
thrW CGU
trpT CCA
tyrT GUA
tyrU GUA
tyrV GUA
valT UAC
valU UAC
valV GAC
valW GAC
valX UAC
valY UAC
valZ UAC
0 5 10
log2FC, 3' end signal
0 5 10
log2FC, 5' end signal
D
tRNA-Gly (UCC)
3'
5'
5' end Fold Change
14
12
10
8
6
4
2
E
B14T: WT Mut
Yin1: WT Mut WT Mut
Time(min): 0 10 0 10 0 10 0 10
cleaved msDNA
msDNA (70nt)
msrRNA (36nt)
TBE-Urea (15%)
GelRed

**Fig S9.- tRNA cleavage analysis by activated retrons**

**A)** Urea-PAGE analysis of total RNA extracted from cells harbouring Sen3 or an empty vector control at various time points following B14T induction. The gel is representative of three independent experiments. **B)** Fraction of internal 5′ ends across different annotated RNA types in selected samples from the experiment shown in Figure 5B, corresponding to the 3′-end analysis shown in Figure 5C. **C)** Enlarged view of the heatmaps shown in Figure 5D, displaying position-specific 3′-end and 5′-end CPM signals for individual annotated *E. coli* tRNAs. Sequences include the anticodon and seven nucleotides upstream and downstream. **D)** Secondary structure of tRNA-Gly(UCC), with nucleotides coloured according to the fold change in 5′-end CPM signal between Sen3-B14T and Sen3 HEPN-inactive mutant-B14T samples, corresponding to the 3′-end analysis shown in Figure 5F. **E)** Control gel showing cleavage of msDNA by B14T. The same gel as in Figure 5G was stained with GelRed. The positions of msDNA and msRNA are indicated. The gel is representative of two independent experiments.
