## Supplementary material for "A DNA-scaffolded Retron Ring Mediates Antiphage Immunity": Table 2

**Cryo-EM data collection, refinement and validation statistics**

|  | #1 Retron Yin1 disrupted rings  (EMDB-58249)  (PDB 31AZ) | | #2 Retron Yin1  full rings  (EMDB-58248) |
| --- | --- | --- | --- |
| **Data collection and processing** |  | |  |
| Magnification | | x165.000 | x165.000 |
| Voltage (kV) | | 300 | 300 |
| Electron exposure (e–/Å^2^) | | ~40 | ~40 |
| Defocus range (μm) | | -0.6, -1.8 | -0.6, -1.8 |
| Pixel size (Å) | | 0.728 | 0.728 |
| Symmetry imposed | | none | none |
| Initial particle images (no.) | | ~13,400,000 | ~340,000 |
| Final particle images (no.) | | 525,790 | 95,927 |
| Map resolution (Å)  FSC threshold | | 3.24  0.143 | 7.20 |
| Map resolution range (Å) | | 2.71->7 | 5.57->19 |
| **Refinement** | |  |  |
| Initial model used (PDB code) | | AF3 models | - |
| Model resolution (Å)  FSC threshold | | 3.20  0.143 | - |
| Model resolution range (Å) | | 2.48->10 | - |
| Map sharpening *B* factor (Å^2^) | | -138.72 | - |
| Model composition  Non-hydrogen atoms  Protein residues  Nucleotides | | 56,198  29,734  2,606  424 | -  -  -  - |
| *B* factors (Å^2^)  Protein  DNA/RNA | | 209.25  273.47 | -  - |
| R.m.s. deviations  Bond lengths (Å)  Bond angles (°) | | 0.004  0.821 | -  - |
| Validation  MolProbity score  Clashscore  Poor rotamers (%) | | 1.26  3.06  0.04 | -  -  - |
| Ramachandran plot  Favored (%)  Allowed (%)  Disallowed (%) | | 97.06  2.94  0 | -  -  - |
| Map-model correlation coefficients  CCmask  CCbox | | 0.7940  0.8916 | -  - |
